# Event Driven Neural Network on a Mixed Signal Neuromorphic Processor for EEG Based Epileptic Seizure Detection

**DOI:** 10.1101/2024.05.22.595225

**Authors:** Jim Bartels, Olympia Gallou, Hiroyuki Ito, Matthew Cook, Johannes Sarnthein, Giacomo Indiveri, Saptarshi Ghosh

## Abstract

Long-term monitoring of biomedical signals is essential for the modern clinical management of neurological conditions such as epilepsy. However, developing wearable systems that are able to monitor, analyze, and detect epileptic seizures with long-lasting operation times using current technologies is still an open challenge. Brain-inspired spiking neural networks (SNNs) represent a promising signal processing and computing framework as they can be deployed on ultra-low power neuromorphic computing systems, for this purpose. Here, we introduce a novel SNN architecture, co-designed and validated on a mixed-signal neuromorphic chip, that shows potential for always-on monitoring of epileptic activity. We demonstrate how the hardware implementation of this SNN captures the phenomenon of partial synchronization within neural activity during seizure periods. We assess the network using a full-custom asynchronous mixed-signal neuromorphic platform, processing analog signals in real-time from an Electroencephalographic (EEG) seizure dataset. The neuromorphic chip comprises an analog front-end (AFE) signal conditioning stage and an asynchronous delta modulation (ADM) circuit directly integrated on the same die, which can produce the stream of spikes as input to the SNN, directly from the analog EEG signals. We show a linear classifier in a post processing stage that is sufficient to reliably classify and detect seizures, from the local features extracted by the SNN, indicating the feasibility of full on-chip seizure monitoring in the future. This research marks a significant advancement toward developing embedded intelligent “wear and forget” units for resource-constrained environments. These units could autonomously detect and log relevant EEG events of interest in out-of-hospital environments, offering new possibilities for patient care and management of neurological disorders.

## 1 Introduction

Epilepsy is one of the most common neurological disorders in the world, affecting nearly 1% of the population worldwide in all age groups [1, 2]. An epileptic seizure is commonly understood as a clinical manifestation of excessive excitation of neurons in the cortex, which can be characterized by locally synchronous high amplitude neural recordings. The research on the detection of epileptic seizures by monitoring the electrical activities of the brain, is a long-standing and massively wide-ranging field of scientific interest [3]. Time series of EEG recordings captured in clinical settings have been generally used as the key signals for this purpose with a high level of confidence [4, 5]. However, the common approach to seizure monitoring currently utilizes a combination of limited in-person clinical observation and a patient-maintained log known as a “seizure diary” outside the clinical setting, which inadvertently suffers from over- and under-reporting [6, 7]. Therefore, long-term always-on monitoring of biosignals outside the clinical environment is a crucial challenge for enabling the treatment of Epilepsy.

Amongst the ongoing revolution in the field of Artificial Intelligence (AI) applied to biomedical applications and biomedical electronics [8, 9], several devices for automated seizure detection have been reported [10]. These devices rely on current advances in biosignal sensing, intelligent Application Specific Integrated Circuits (ASICs), and machine learning algorithms [8, 11, 12, 13]. While most record multiple biopotentials, such as EEG, electrocardiogram, electromyography, electrodermal activity, and accelerometry, the EEG signal remains the most reliable and gold standard. Two pioneering devices in the domain of EEG-based monitoring are the “Seizure Advisory System” (SAS) developed by NeuroVista Corporation [14] and the “Responsive Neurostimulation System” (RNS) of Neuropace Corporation [15]. More recently, dedicated digital accelerators such as BioGAP [16] have also been proposed to tackle the problem of wearable biosignal monitoring [17, 18]. Although promising, such digital systems, based on general-purpose programmable computer architectures, suffer from the same limitations of conventional digital computing approaches in terms of power consumption for long-term beyond-laboratory monitoring of temporally slow and sparse signals [19].

Conversely, analog neuron and synapse circuits that use the physics of the transistors to emulate the biophysics of biological neural circuits [20, 21] represents a promising approach for implementing ultra low-power always-on electronic processing systems aimed at biomedical signal processing [22, 23]. Mixed-signal analog/digital neuromorphic computing systems with analog neural and synaptic *compute* and asynchronous on-demand digital routing operate very efficiently in resource-constrained environments, enabling long-term monitoring of biosignals and potentially leading to the design of wearable seizure detection units (Figs. 1a and c).

**Figure 1.**
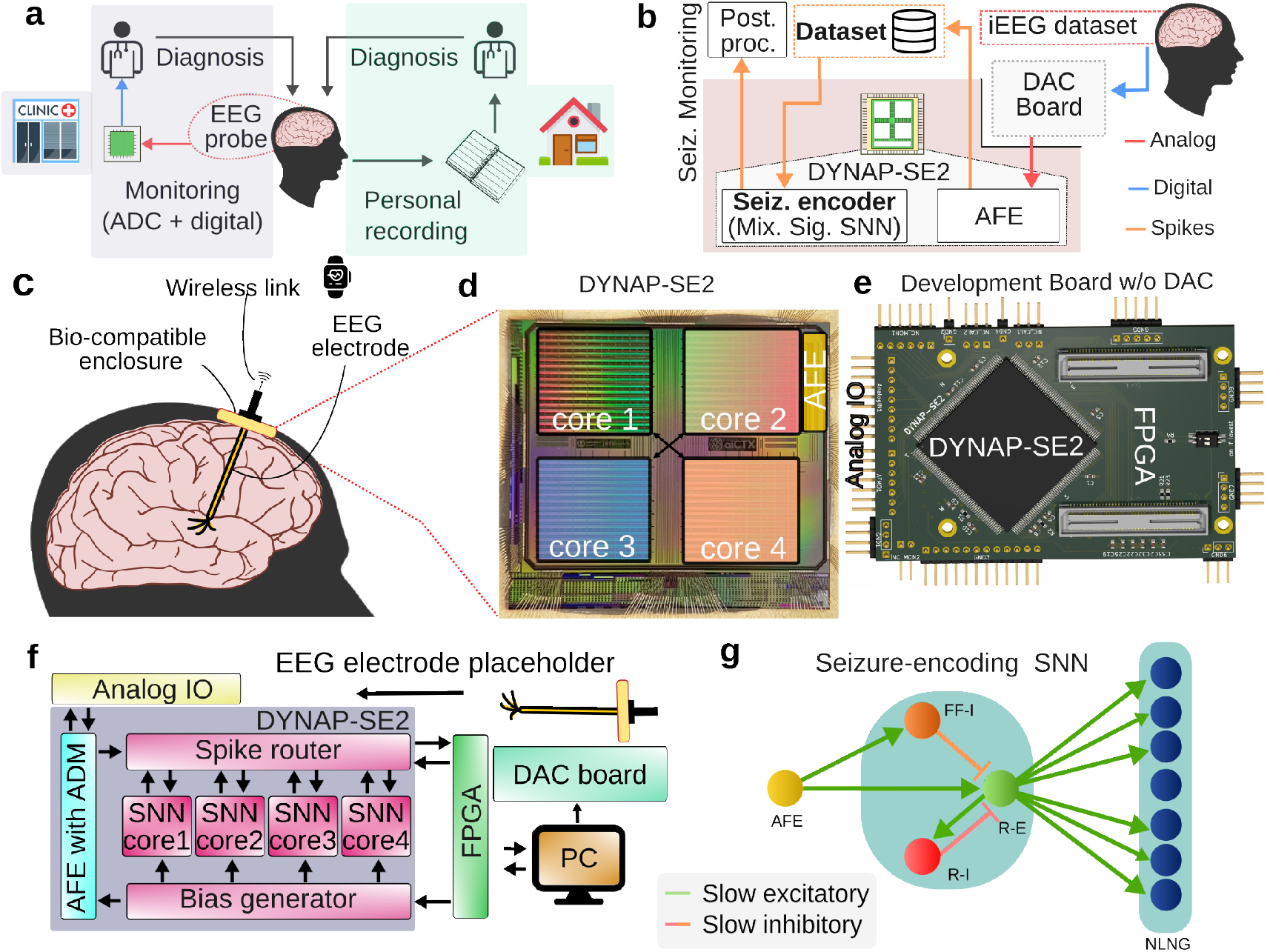
Overview of seizure detection process, and of the setup constructed for this work; **a** SOA (state of art) approaches for seizure detection and diagnosis; personal monitoring with a seizure diary (right side), inpatient seizure detection using Analog-to-Digital Converters (ADC) and digital processing (left side); **b** Proposed system; seizure encoding on DYNAP-SE2 with iEEG signals using on-chip AFE and SNN; **c** Envisioned system; real-time seizure detection with industry-standard EEG electrodes using an SNN on a multi-core neuromorphic processor within a bio-compatible enclosure; **d** DYNAP-SE2 die with SNN cores and AFE; **e** Development board for DYNAP-SE2; **f** Block diagram of the boards and DYNAP-SE2: an FPGA controller, DYNAP-SE2 board, Digital-to-Analog (DAC) converter mimicking the EEG electrode, interfacing with the AFE; the FPGA sends input spikes to the spike cores and configures the AFE and SNN cores; a DAC board sends analog EEG signals to AFE; **g** proposed SNN architecture with the Analog Front-End (AFE) input, Recurrent Excitatory/Inhibitory (R-E/I) and Fast-Firing Inhibitory neuron (FF-I) neurons of translation layer and neurons of Non-Local Non-Global (NLNG) layer for seizure detection.

However, unlike digital AI accelerators, these systems cannot be “programmed” to solve specific tasks using traditionally-utilized machine learning algorithms. The primary challenge is to establish methods for designing SNNs under the constraints of the hardware, i.e., a fixed number of neurons and synapses, limited precision, variability in the network parameters, and sensitivity to noise, that can solve the problem at hand robustly and accurately by leveraging the inherent dynamics of analog synapses and neurons. Indeed, the intrinsic variability and mismatch of mixed-signal systems, coupled with event pulse-based operations makes programming a noise-resistant and robust SNN model onto such devices a non-trivial and complex undertaking [24].

This paper addresses this challenge and proposes a novel two-layered SNN compatible with mixed-signal analog/digital neuromorphic hardware that amplifies and extracts the intrinsic partial synchronization (i.e., *Chimera*) present but not prominent in input signals during seizure periods [25]. We validate this SNN on a full-custom neuromorphic hardware pipeline (Fig. 1b) that does not require any digital compute element in the loop. Moreover, we demonstrate the proper operation of this SNN hardware implementation on a DYnamic Neuromorphic Asynchronous Processor (DYNAP-SE2) [26] (Figs. 1d and e) with an estimated power consumption of 150 *μW*. The proposed SNN is both biologically inspired and (as a consequence) compatible with neuromorphic hardware.

By designing a hardware-aware shallow SNN that couples the sparsely encoded input spike-trains in a specific network topology with adjacent but non-global connections (Fig. 1g), the network amplifies correlations and forms localized synchronized clusters during seizure (ictal) periods, mirroring partial synchronous states observed in the EEG recordings during seizures [27, 28]. As a major advance over previously reported related approaches [29, 30], signal conditioning, spike encoding, and SNN processing are all realized directly in the neuromorphic hardware and on the same chip (Fig. 1e). Although classification using the SNN output spikes is performed off-chip with a linear classifier, this work represents the first completely integrated proof-of-concept towards the encoding of seizures through partial synchronization in real-time using an ultra-low power neuromorphic computing substrate, providing a foundation for future always-on seizure monitoring devices.

## 2 Results

The proposed SNN encodes the partial synchronization of brain states during seizure (ictal) periods by means of firing rate-based filtering and a specific connectivity structure (Fig. 1g). As an alternative to the common approach in biosignal monitoring, involving short in-patient sessions and self-maintained records (Fig. 1a), the real-time on-chip seizure encoding pipeline of fig. 1b, uses event-encoded EEG signals as input.

In the subsequent sections, we first describe the AFE circuit that contains low-noise amplifiers, filters, and the ADM circuit [31] directly integrated into the DYNAP-SE2 chip. This AFE is employed to convert the EEG signals from a publicly available iEEG dataset [32] into streams of spikes (events). This is followed by details of the implemented SNN architecture and an explanation of the operation of the pipeline in encoding seizures. Afterward, we utilize a linear classifier in the post-processing step to benchmark the extracted features of the SNN and show the classification of seizure periods with high accuracy and specificity.

### 2.1 Asynchronous Delta Modulation (ADM) of EEG signals

The first stage of the processing pipeline (outlined in Fig. 1f)is comprised of the AFE and ADM blocks embedded in the DYNAP-SE2 (Figs. 1b and d), for converting analog EEG signals into streams of digital events. Figure 2 shows recorded analog intracranial EEG (iEEG) waveforms from two input channels (Channel 12 and 13 of a single seizure of patient 1), selected from the SWEC dataset [32]. These traces were obtained by converting the digitized SWEC dataset waveforms back into analog signals using a DAC setup (Fig. 1f) and amplified by the initial stage of the AFE. The amplified signal stream of the AFE is then inputted to the ADM, which produces asynchronous streams of pulses marked Up and Down, depending on the polarity of the slope of the analog waveform (See Methods section for details). The Up and Down events are shown above and below the analog waveforms and the event rate is color-coded as per the colorbar (Fig. 2). A close-up of 16 seconds of the first channel (zoomed plot above Fig. 2a) shows the increase of event rates and their burstiness during the seizure (ictal) periods. Figure 6b presents the oscilloscope traces of the analog EEG signal. The conversions with the AFE are recorded in an “event-based hardware encoded dataset” (see data availability). In total, we encoded and made publicly available [33] data for ten seizures, two per patient for a total of five patients. The events from the ADM circuit for all channels are fed synchronously with a single pulse train per channel (due to the limited fan-in of the neurons) into the SNN core of the DYNAP-SE2 (See Methods).

**Figure 2.**
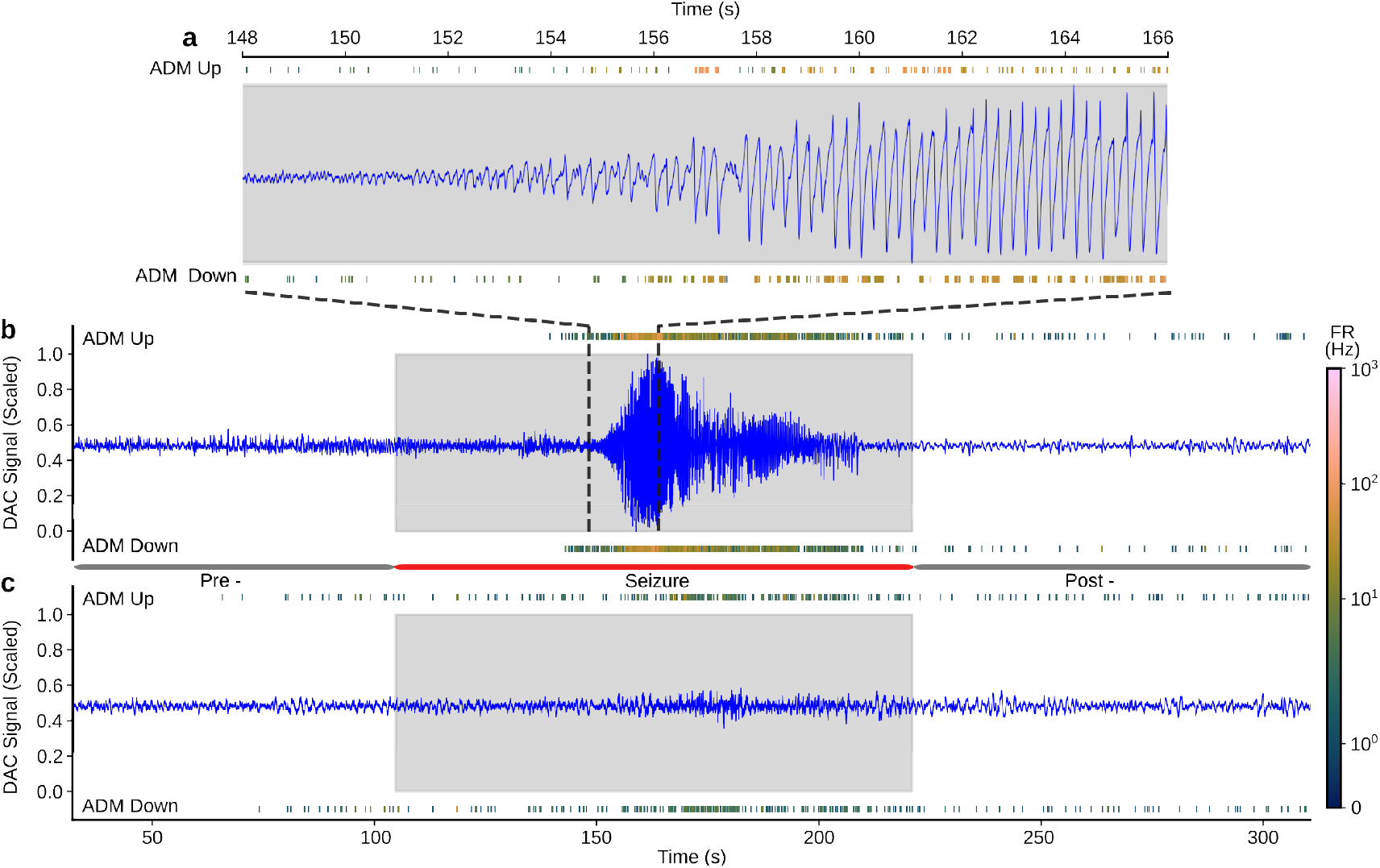
Snippets presenting 320-second long DAC converted analog iEEG signals from two channels and the encoded events from the ADM (*UP* and *DOWN*) on DYNAP-SE2. The event rate (above and below the DAC-generated signal) is encoded in color with the seizure period denoted by the grey background; **a** Analog waveform of one iEEG channel and ADM events (with inset above showing a close-up of 16 seconds); **b** Analog waveform of another iEEG channel and ADM events.

### 2.2 The SNN model for firing-rate filtering and encoding correlation

The second stage of the pipeline outlined in Fig. 1f, is the SNN model implemented on the DYNAP-SE2. Figure 3a shows the SNN architecture. It comprises two layers: a “translation” layer and a “non-local non-global” (NLNG) encoding layer. The translation layer, adapted from Sava et al. [34], consists of Recurrent Excitatory (R-E) and Inhibitory (R-I) neurons, combined with a Fast-Firing Inhibitory neuron (FF-I) depicted in Fig. 3a (left). The FF-I and the R-E neurons receive events from the ADM-converted event streams for iEEG signal channels via a slow-excitatory synapse. Further, the R-E neurons are inhibited by the output spikes from FF-I neurons via a slow-inhibitory synapse. The R-E and R-I neurons represent a neuron pair that both excite and inhibit each other, constructing an excitatory-inhibitory (E-I) balance pair.

**Figure 3.**
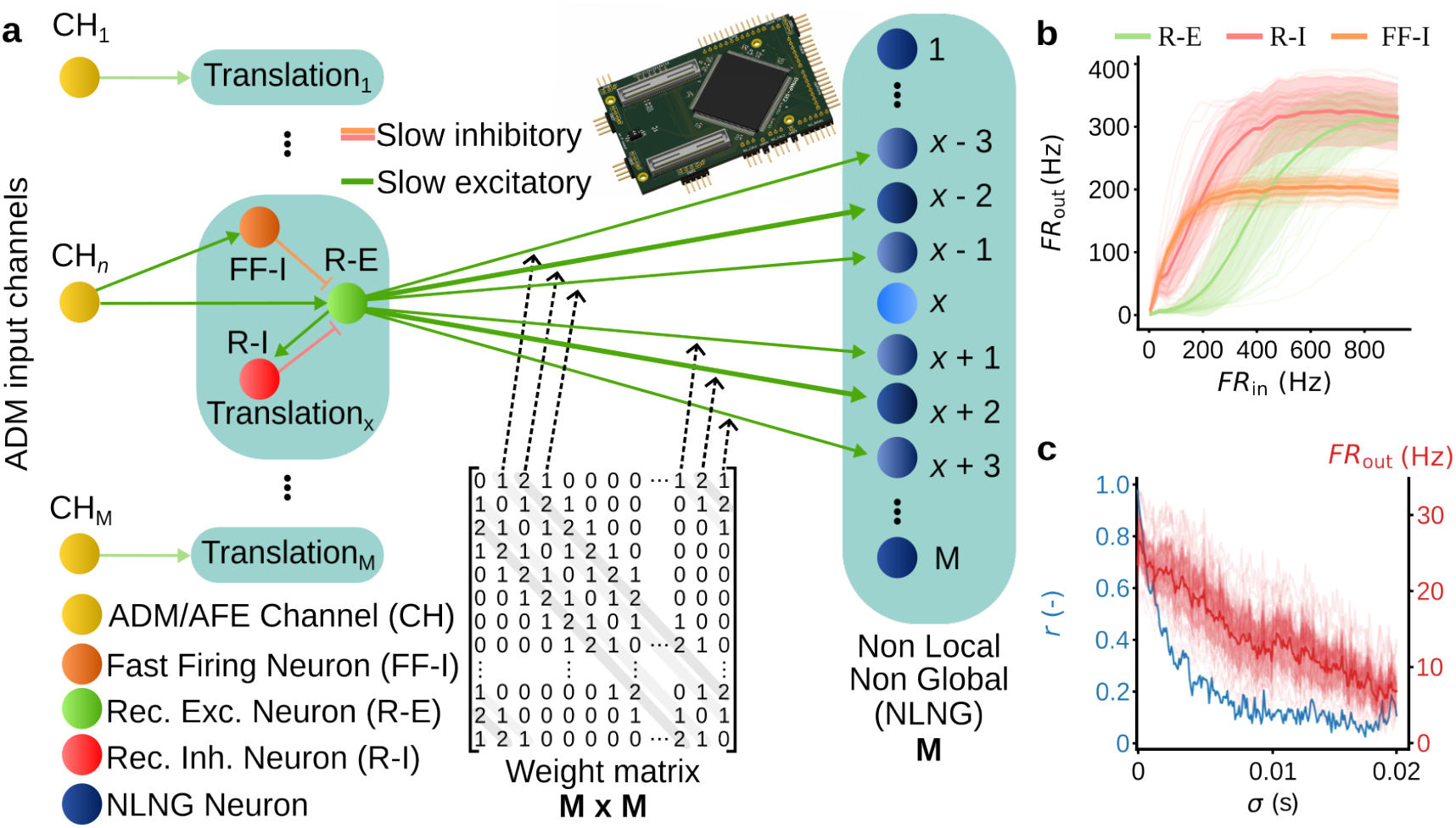
**a** Structure of the on-chip SNN; a translation layer containing 3 types of neurons, Fast-Firing Inhibitory (FF-I), Recurrent-Excitatory (R-E), and Recurrent Inhibitory (R-I) neurons. This layer performs firing-rate-based filtering, followed by a Non-Local Non-Global (NLNG) layer. Synapse types slow excitatory (NMDA like) and slow inhibitory (GABA-B like) are color-coded; Weighted connectivity matrix display projections from R-E to NLNG neurons with a spatial 1 – 2 – 1 (mirrored) connections **b** Firing rate input-output curve for different types of neurons namely FF-I (orange),R-E (green) and R-I (red); color coded by type; as a result of Poisson-generated input spikes to the translation layer; **c** In blue, the mean Pearson correlation coefficient of M = 42 input channels over the standard deviation of jittering in spike times at input firing rates of 50 Hz. In red, the output firing rate of the non-local non-global encoder as a result of the jittered input spike trains

Figure 3b shows the firing rate input-output curve of the translation layer in the DYNAP-SE2 based on input Poisson trains of varying firing rates. The synaptic parameters for synapses attached to different neuron types (R-E, R-I, and FF-I) have been set heuristically to achieve the frequency transfer curve in Fig. 3b. All other parameters including the weights were left constant (see Methods for parametric values). The slow excitatory synapses attached to FF-I neurons have a high bias current associated with its synaptic time constant (resulting in a slower decay as compared to R-E/I), leading to a firing rate that saturates at around 180 Hz making inhibition to R-E constant for input firing rates >200 Hz (responsible for the lower bound demonstrated in Fig. 3b). The slow inhibition of the R-E neurons by R-I neurons results in a near-linear output firing rate for input firing rates between 200 Hz to 600 Hz. These neurons inhibit both very high and low-frequency input spikes. Thus, the above described three neuron blocks act as a spike-based filter, i.e., the excitation of the R-E neuron is highly sensitive to the 200–600 Hz firing rate band while insensitive to inputs from outside this band. Further, the block linearly maps the firing rates of the ADM/AFE event streams to increase the dynamics range [34] of the input to the next layer (Fig. 3b).

The second layer, termed the NLNG encoding layer, is excited by the R-E neurons of the translation layer with a specific connectivity structure. The connections consist of slow excitatory synapses with constant synaptic weights. The constant weight per synapse is chosen to make sure that the membrane potential of the neurons reaches close to the set threshold. We used multiple connections per projection from the source to the destination, to change the effective synaptic weight. The connectivity structure is described by a 1 – 2 – 1 – 0 – 1 – 2 – 1 connectivity between the R-E and NLNG neurons (Fig. 3a) where 2 represents two connections with constant weight per projection (see Methods Sec. 5.4 for parametric values).Choice of such constant weight assignments for synapses stem from inherent hardware constraints including low bit resolution in weight assignments, limitation for setting parameters and inherent variability of analog synapse circuits (see Methods Sec. 5.4). To give an example, we consider M input neurons (encoding the M EEG channels, for the case of a particular patient), producing M outputs from R-E neurons as per the frequency response curve shown in Fig. 3b. Here, the R-E neuron from a translation block x is connected to the neighboring NLNG neurons (x-3,x-2,x-1 and x+1,x+2,x+3, mirrored) with 1 – 2 – 1 connectivity depicting one connection, two connections with constant weight (connecting same pre- and post- neuron), and one connection per projection, respectively. This connectivity pattern is mirrored on both sides while ignoring NLNG neuron x itself. The weighted connectivity matrix is shown in Fig. 3a, bottom and, is similar to a regular graph [35]. Such connectivity patterns are established for each R-E neuron connected to the NLNG layer with closed boundary conditions.

This connectivity matrix between these layers is determined by a Evolutionary Neural Architecture Search (ENAS) [36]. Utilising a differential evolutionary algorithm with a cost function defined as the maximum separation between output firing rate from non-correlated input and correlated Poisson input, the ENAS approach leads to the unique non-local non global structure with the 1 – 2 – 1, mirrored connection. Figure 3c shows an inverse relationship in the output firing rate of the NLNG layer from DYNAP-SE2 with the correlation between input spiketrains that approaches linearity. These spike trains are generated from a Poisson process and iteratively jittered with jitter noise taken from a normal distribution and input simultaneously across M channels to the chip. The NLNG layer exhibits a firing rate near >30 Hz for fully correlated input spike trains, decreasing to 20 Hz and 10 Hz for less correlated spike trains, with mean Pearson correlation coefficient of *r* =0.255 and *r* = 0.114, arising from spike time jitter noise at standard deviations of *σ* = 0.005 s and *σ* = 0.015 s respectively (Fig. 3c). This experiment shows the functionality of this ENAS proposed layer and its ability to encode correlation between input channels by means of firing rate for ideal Poisson inputs.

### 2.3 Encoding seizures with partial synchronization using the hardware aware SNN

The SNN network presented in the previous section forms the core of our pipeline, which is used to encode and extract features based on correlated inputs, namely the event-encoded EEG signals. As the dynamics of these events are substantially distinct from Poisson-based spike trains (containing bursty spikes as shown in Fig. 2a, top), we reduced the bias current for synaptic time-constants (leading to a faster decay as compared to R-E/I) for synapses attached to the NLNG neurons compared to the previous section. We further analyzed the output of the NLNG neurons by means of synchronization matrices.

Figure 4a shows an example of 42 raw iEEG waveforms of the first seizure of patient 1 from the SWEC dataset with annotated seizure (ictal) period. The events of each EEG channel are simultaneously fed to the SNN cores of DYNAP-SE2 in real-time (Fig. 4b, row-1). The translation layer rejects the lowest firing rate input spikes and reduces high input firing rates (Fig. 4b, rows 2,3,4) while keeping a linear relationship between the input and output firing rates in intermediate areas similar to Fig. 3b.

**Figure 4.**
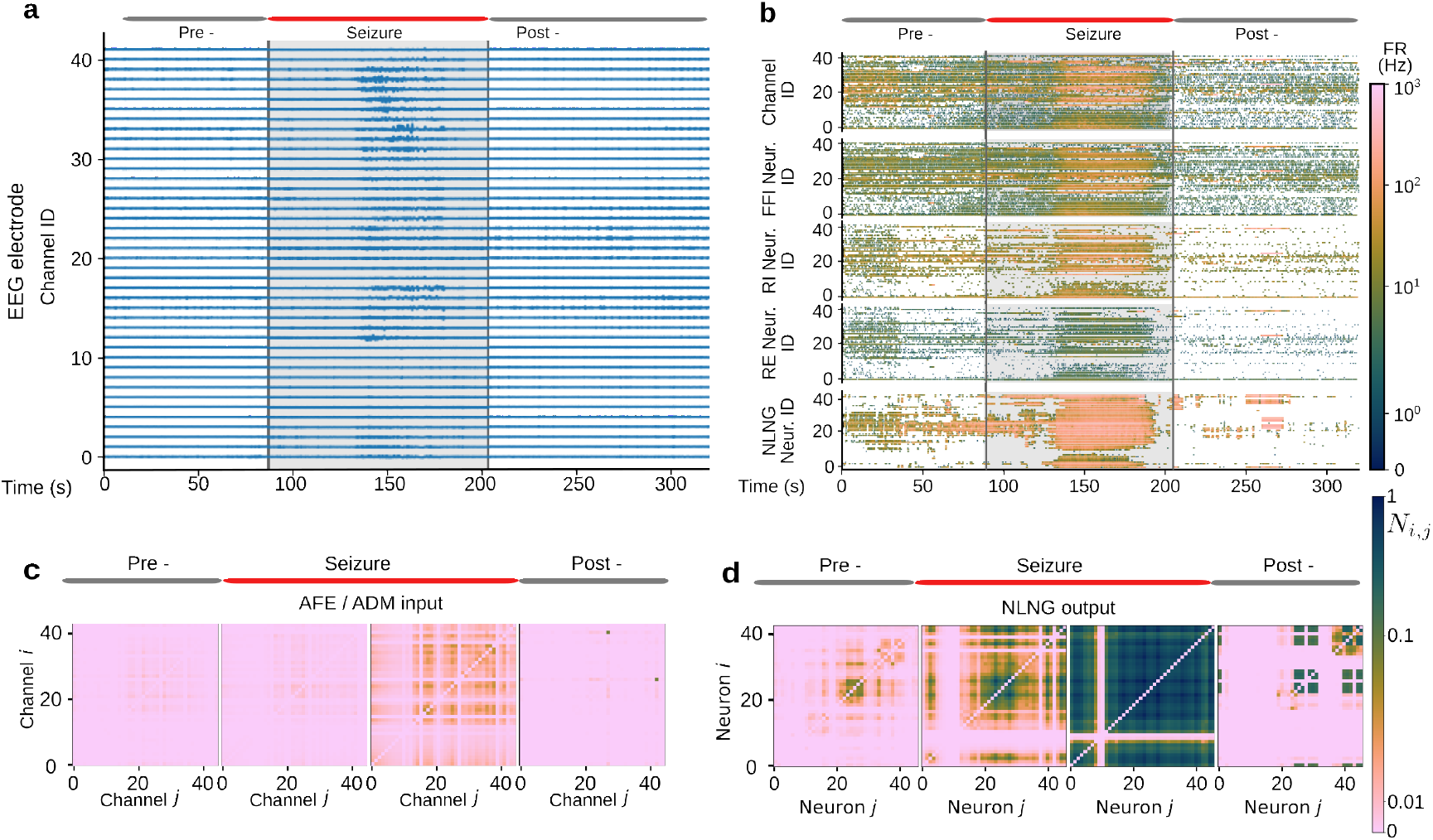
**a** 42 channels of input EEG signals from patient 1, seizure 1; **b** Output rasterplots of the DYNAP-SE2 on-chip implemented network. Top row to bottom: ADM-encoded input of **a**, FF-I neurons’ outputs, R-I and R-E (E-I balance pairs) neurons’ outputs, and outputs of the NLNG neurons. The firing rate is color-encoded per one-second time-segment; Normalized synchronization matrices between channel and neuron pairs of **c** AFE input and **d** NLNG output, respectively. From left to right: the pre-ictal, ictal (seizure), and post-ictal periods. The synchronization matrices during the ictal period (2nd and 3rd column) are created from the first half and second half of the seizure in time, respectively. Note the logarithmic scale in color-bar, used to highlight minute correlations in ADM/AFE synchronization matrices, which reduces the dynamic range of NLNG synchronization matrices.

The NLNG layer shows a high firing rate for the R-E neuron inputs that is correlated locally, (see 125 s <t< 190 s in Fig. 4b). In the pre-seizure period, the NLNG layer does not spike at firing rates >100 Hz. During the seizure (as indicated in Fig. 4b), the firing rate of a group of neurons in the NLNG layer increases substantially to values above 100 Hz and continues to increase until the seizure ends. Such mode of operations across different SNN blocks holds across all the patient data analyzed here (See supplementary for similar raster plots of SNN blocks for all seizures of all patients). For these tasks, the power consumption of the neurons in the SNN to produce the output spike rates at the standard 1.8 V supply voltage was estimated to be 150 *μW* (with 2.8 *μW* power per input channel) on average across all patients (See Methods for details about the calculation of the SNN power consumption).

In addition, we have employed kernelized spike trains with causal exponentially decaying functions [37], denoted as *f*_*i*_ (*t*) where *i* indicates the NLNG neuron index. Using these kernelized time-continuous spike-trains, we determine a measure of synchronization by taking the Manhattan (*L*_1_) norm of the inner product between neuron pairs *i* and *j*, *N*_*i*, *j*_ = ∥ < *f*_*i*_, *f* _*j*_ > ∥ _1_ and average them over a fixed time period; see Methods for a step by step explanation. In essence, we define synchronization as the similarity of spike-trains between neurons and quantify it with the time averaged norm *N*_*i*, *j*_ .

For the same example, figure 4 (in bottom) shows the normalized synchronization matrices for all pairs of AFE event streams/channels (Fig. 4c) and all NLNG neuron *N*_*i*, *j*_ pairs (Fig. 4d) for a single seizure. Choosing four time windows, namely, pre-seizure, first and second half of the seizure and post-seizure, we computed the norm *N*_*i*, *j*_ for all pairs and averaged them over the above mentioned time-windows. Figure 4d clearly shows formation of locally synchronised clusters during the seizure time-window. Comparing figures 4c and d, the matrices demonstrate the enhancement of the inherent correlation structure present when seizures occur (See supplementary for similar matrix plots of AFE and NLNG for all seizures of all patients).

Figure. 5 shows the normalized synchronization matrices for the NLNG layer (where the norm *N*_*i*, *j*_ is averaged over the entire seizure period) for all seizures of all patients analysed here. A distinct cluster formations, characteristics of each patients can be observed in the columns of Figure. 5. Additionally, the seizures from the same patient displayed similar cluster formations across top and bottom rows of Figure. 5. These observations confirm that these emergent synchronization phenomena are replicated by the NLNG layer across all patients during the seizure period. The clusters emerging in Fig. 5 for different patients in the seizure period show emergent localized synchronization holding true for all patients.

**Figure 5.**
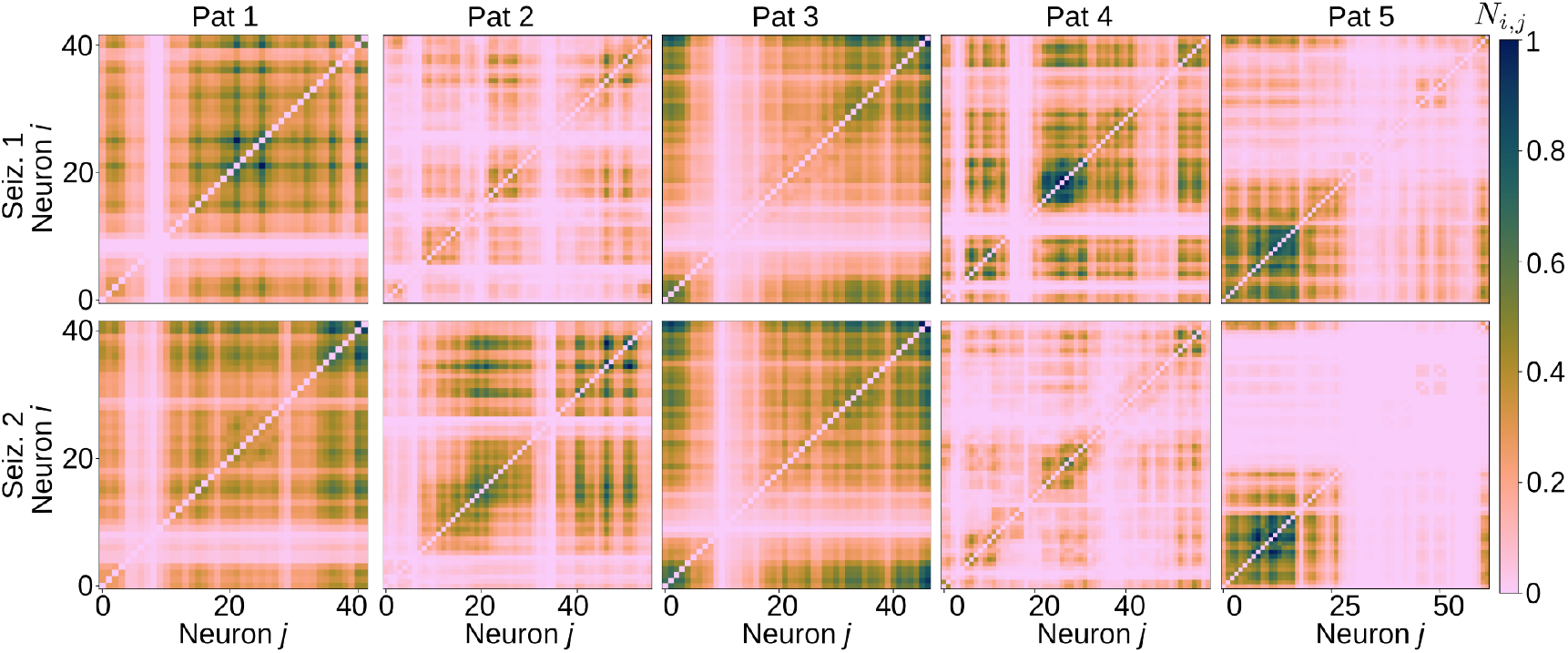
The synchronization matrices for all pairs of neurons in the NLNG layer during the seizure period; Note two seizures are analysed for each patient and they correspond to top and bottom row, respectively. Synchronization matrices are presented for five patients (as shown in column headings). Note the linear scale in color-bar, used to increase the dynamic range of NLNG sync. matrices.

**Figure 6.**
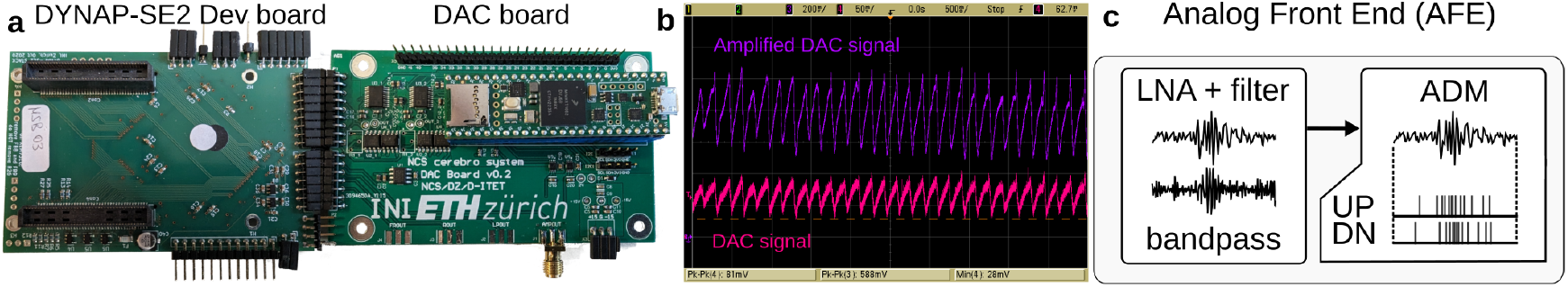
The utilised setup and AFE; **a** Photograph of the boards, an FPGA board, DYNAP-SE2 board with DYNAP-SE2 chip on the underside, DAC converter board with a Teensy microcontroller for interfacing with the AFE **b** Oscilloscope traces depicting the incoming DAC-generated signal (trace 4, in pink) and the AFE amplified signal (trace 3, in purple); **c** Internal components of the AFE, a Low-Noise Amplifier (LNA) with bandpass filter-bank, followed by the ADM.

### 2.4 Detecting seizures using SNN encoded features

The SNN on DYNAP-SE2, illustrated in Fig. 3, functions as a feature encoder, transforming ADM-encoded data from M independent input channels into a richer representation, where the NLNG layer captures complex temporal-spatial dependencies between channels, facilitating seizure detection. As a post-processing step, we explored how well the SNN-encoded features could perform with a conventional and widely used machine learning classifier. The output spike trains from the NLNG layer and the original input from the AFE/ADM were employed as features, demonstrating the SNN’s capacity on DYNAP-SE2 to differentiate ictal from interictal activity by using Linear Support Vector Machines (LSVM).

We investigated two distinct spike-driven feature extraction approaches: firing rates and synchronization matrices derived from the spikes in the AFE/ADM and NLNG layer. The NLNG and AFE spike trains are binned into non-overlapping one-second windows, channel-by-channel, normalized, and collectively referred to as firing rates (FR). As a second feature extraction approach, we investigated synchronization matrices, by transforming spike trains into continuous functions through kernelization as described in detail in the Methods section. Synchronization between pairs of neurons is quantified using the inner product of their continuous spike trains, followed by calculating the Manhattan (*L*_1_) norm of the inner product to capture the temporal alignment of spikes across channels. These matrices are calculated over sliding windows (window size = 1 sec, no overlap), where each *M* × *M* synchronization matrix is symmetric (See Methods Sec. 5.6.2). To avoid redundancy, only the upper triangular part (excluding the diagonal) is extracted, flattened into a 1D feature vector of dimension 1 × *M* (*M* − 1) /2, and normalized to ensure that they are on a comparable scale before being used as feature vectors for SVM classification. The spike trains and features are publicly available (see data availability statement at Sec. 6.4) for reproducibility and further study without requiring access to the DYNAP-SE2 hardware.

By adopting a patient-specific training and testing approach, we aimed to mitigate the challenges posed by variability in the number of channels and duration of seizure recordings between patients, which complicates cross-patient generalization. This decision also reduces the risk of overfitting, as it ensures that the models are tailored to the unique characteristics of each patient’s seizure data. Furthermore, for a more balanced and robust evaluation, we applied stratified 5-fold cross-validation for each patient, where each fold maintains an equal representation of seizure and non-seizure windows. Bayesian optimization was used to fine-tune the hyperparameters of SVM classifiers.

Table 1 describes the LSVM results for seizure detection per patient based on the ADM and NLNG synchronization measure and Table 2 based on the firing rates. The classification metrics of interest are accuracy, sensitivity, specificity, F1-score, and Area Under the Curve (AUC). The variability in patient-specific seizure characteristics is reflected in the classification results. Overall, NLNG FR slightly outperforms ADM FR in most metrics. NLNG FR is more effective at detecting seizure windows, evidenced by its superior average AUC 80.91% compared to 78.58% and also shows higher accuracy (78.35% vs. 77.22%), sensitivity (66.71% vs. 58.56%), and F1-score (69.96% vs. 65.13%) on average across patients. However, average specificity is slightly higher for ADM FR (89.55% vs. 85.54%), indicating that it performs better at correctly classifying non-seizure windows. The results in Table 1 indicate that both ADM and NLNG synchronization features yield comparable performance, when using the same sliding window technique (1 sec with no overlap) and linear kernel. However, there is a noticeable decrease in the average performance across patients compared to the firing rates. Further comparative analysis investigating the influence of nonlinear SVM with RBF kernel and other hyperparameters considering temporal dependencies (window size, step size) is reported in the Supplementary Material.

**Table 1.** Seizure classification performance using LSVM with ADM/AFE and NLNG synchronization features across patients.

| Metric | Feature | ID1 | ID2 | ID3 | ID4 | ID5 | Avg. |
| --- | --- | --- | --- | --- | --- | --- | --- |
| Accuracy [%] | ADM SYN | 74.70 | 53.54 | 75.54 | 81.76 | 59.97 | 69.10 ± 10.57 |
| | NLNG SYN | 78.78 | 69.76 | 70.05 | 76.45 | 60.35 | 71.07 ± 6.42 |
| Sensitivity [%] | ADM SYN | 76.77 | 30.22 | 66.69 | 62.09 | 58.60 | 58.87 ± 15.57 |
| | NLNG SYN | 78.92 | 58.11 | 66.80 | 55.87 | 58.60 | 63.56 ± 8.48 |
| Specificity [%] | ADM SYN | 72.19 | 70.71 | 80.54 | 89.53 | 61.68 | 74.93 ± 9.43 |
| | NLNG SYN | 78.60 | 84.17 | 71.94 | 88.33 | 62.49 | 77.10 ± 9.14 |
| F1-Score [%] | ADM SYN | 76.68 | 32.33 | 65.06 | 64.97 | 60.29 | 59.86 ± 14.79 |
| | NLNG SYN | 80.87 | 63.52 | 60.07 | 61.16 | 60.91 | 65.3 ± 7.86 |
| AUC [%] | ADM SYN | 82.42 | 45.69 | 80.80 | 77.93 | 61.48 | 69.65 ± 14.12 |
| | NLNG SYN | 84.58 | 65.72 | 78.47 | 73.53 | 62.71 | 73.80 ± 8.72 |

**Table 2.** Seizure classification performance using LSVM with ADM/AFE and SNN-Encoded NLNG features across patients.

| Metric | Feature | ID1 | ID2 | ID3 | ID4 | ID5 | Avg |
| --- | --- | --- | --- | --- | --- | --- | --- |
| Accuracy [%] | ADM FR | 80.33 | 72.64 | 78.87 | 90.09 | 64.34 | 77.22 ± 11.32 |
| | NLNG FR | 81.80 | 77.40 | 78.15 | 86.04 | 68.02 | 78.35 ± 10.95 |
| Sensitivity [%] | ADM FR | 68.90 | 34.07 | 65.21 | 66.00 | 63.00 | 58.56 ± 22.99 |
| | NLNG FR | 76.67 | 53.18 | 78.35 | 60.67 | 67.49 | 66.71 ± 21.08 |
| Specificity [%] | ADM FR | 94.00 | 99.69 | 86.70 | 99.24 | 65.88 | 89.55 ± 8.95 |
| | NLNG FR | 87.82 | 94.38 | 78.02 | 95.68 | 68.63 | 85.54 ± 12.65 |
| F1-Score [%] | ADM FR | 77.85 | 42.79 | 66.83 | 74.75 | 65.19 | 65.13 ± 21.36 |
| | NLNG FR | 81.72 | 58.94 | 73.32 | 67.40 | 68.94 | 69.96 ± 17.31 |
| AUC [%] | ADM FR | 86.41 | 74.42 | 81.28 | 84.46 | 66.52 | 78.58 ± 15.39 |
| | NLNG FR | 86.80 | 80.82 | 80.21 | 86.06 | 69.97 | 80.91 ± 14.80 |

## 3 Discussion

This work presents a neuromorphic approach towards a real-time seizure monitoring system by employing asynchronous delta modulation based encoding and analog neuronal dynamics on-chip. Instead of the conventional machine learning training methods for solving bio-signal processing tasks with digital computing systems, the approach proposed capitalizes on the rich dynamics arising from sub-threshold analog circuits to produce the desired results. We show how this mixed-signal hardware setup preserves all of the information present in the original data, by comparing the classification accuracy on the spiking data with a Linear Support Vector Machine (LSVM).

Moreover, compared to conventional seizure detection systems [10], the prototype embedded system developed here, does not operate based on a global system clock but processes signals in a real-time data-driven mode with asynchronous encoding, spike routing, and analog neurons [26]. This setup thus by design, foregoes the need for memory buffers (which commonly consume the majority of the power budget on system level) required in clock-based traditional approaches. This is a distinct advantage in terms of power consumption and scalability for always-on applications as for most of its operational time, in the absence of high energy inputs, the system is silent, significantly prolonging the monitoring period [19, 38].

The observation that the NLNG layer imitates the ictal state in real-time by forming partially synchronized clusters within itself can be studied to understand the temporal characteristics of seizure patterns. Future work could involve investigating the formation of the synchronized clusters through time and mapping their relation to the location of the recording channels, helping in tracking the progression of the seizure. We also recognise the role of recording electrode types (strip, grid, or depth type; see sec. 5.7) for each case as a contributing factor for the emergent synchronization clusters. However, unavailability of such meta-information in the public dataset prohibits us from evaluating our assumptions. Regardless, we believe such work can help in the identification of the Seizure Onset Zones (SOZ) (with mapping of such clusters to physical recording locations) that are responsible for seizure generation and provide insights into the disorder itself. In the context of intracranial EEG and epilepsy surgery, this approach could serve as a valuable tool for inferring and localising SOZs in real time in clinical settings and potentially complement existing pre-surgical evaluation techniques to guide surgical interventions.

The correlation patterns between neuron pairs in the AFE event streams and the NLNG spike streams vary significantly (Figs. 4c and d). This suggests a future investigation avenue to design an event-based output layer that have the potential for developing personalised seizure detectors based on the correlation patterns of neuron pairs taking advantage of the emergent but varied, locally synchronized clusters. This approach can also be expanded for the identification of different temporal zones in a seizure event and might help with the personalised classification of seizure types based on partially-synchronized cluster patterns. Classification with a LSVM shows that the NLNG neurons are capable of separating ictal and non-ictal periods. However, we note that the accuracy of such an offline linear classifier is lower in comparison to standard ML approaches and do not significantly change between AFE-encoded event streams and NLNG-encoded spike streams, regardless of features used. This observation, coupled with the limited sample size analyzed in this study and the lack of fine-tuning to patient-specific characteristics, suggests that more tailored approaches or additional features might be required to enhance detection performance. Further research is necessary to develop either on-chip event-based methods or off-chip algorithms for decoding of the NLNG layer into different classes dealing with either seizure types or the characterization of temporal domains. This work serves as a gateway to a more universal approach to exploiting the dynamics of analog neurons and their use in classification tasks. The NLNG layer, coupled with event-encoded input channels through a tuned translation layer, showcases the potential of analog computation in encoding the non-linear dynamics of a complex system. As is exemplified by the structure of the NLNG weight matrix, we believe network structure, emphasizing topology and reciprocity, appears particularly well-suited to the intricacies of mismatched neuromorphic devices. Given the inherent variability of on-chip weights and parameters in mixed-signal or analog neuromorphic devices [39], integrating the lottery ticket hypothesis [40, 41] with the hardware’s dynamical and topological properties could pave the way for additional advancements. We believe this interdisciplinary approach could open up new avenues for the widescale adoption of these devices in a diverse number of applications that are not limited only to bio-medical domains but expand into any complex systems.

## 4 Conclusions

This article presents a first step towards a real-time event-based seizure monitoring system. We introduce an event-based EEG seizure dataset created with an on-chip AFE followed by an SNN with special connectivity to encode seizures with a mixed-signal neuromorphic device. The SNN on-chip shows a proof-of-concept to encode seizures during ictal periods using its inherent dynamical properties.

To our knowledge, this is the first human EEG seizure dataset that has been encoded into events using a realtime delta-modulator. The dataset and synchronization encoded spike trains have been made publicly available [33]. The framework of this investigation provides detailed specifications for the design of new front-ends and SNNs implemented on neuromorphic hardware, that show potential for optimized on-edge seizure monitoring. Although incomplete due to the use of an off-chip linear classifier, we believe this framework represents a foundation towards designing more general class of always-on embedded and low-power neural network computing systems for a wide range of biomedical signal processing tasks.

## 5 Methods

The DYNAP-SE2 neuromorphic processor contains analog exponential integrate and fire (Exp I&F) neuronal and synaptic circuits with spike-frequency adaptation. The DYNAP-SE2 also hosts the AFE including the level-crossing Asynchronous Delta Modulator (ADM) circuit on the same die. Here, we describe the equations of the Exp I&F neurons and synapses derived from the circuit equations of DYNAP-SE2. This is followed by an explanation of the AFE and a description of the neuron cores of DYNAP-SE2. Afterward, we introduce the approach to compute the correlation and construct the synchronization matrices, followed by a detailed event-based dataset description.

### 5.1 Neuron and synapse models

The neurons of the SNN in this work consist of current-mode Exp I&F neuron circuits [26]. The detailed description of the circuit equations in the sub-threshold regime and their deduction leading to this simple form is described by Chicca et al. [21]. Omitting the adaptation term, this equation is shown by

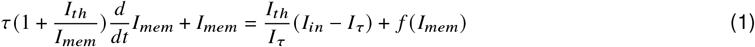

where *I*_*mem*_ is the subthreshold current depicting the membrane potential of the silicon neuron. Other terms including the time constant and the threshold can be defined as follows:

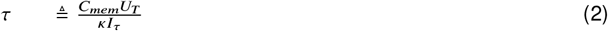

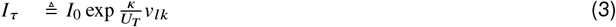

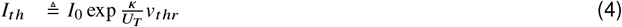

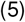

*U*_*T*_ is the thermal voltage, *κ* is the subthreshold slope factor stemming from the capacitive coupling ratio from gate to channel and *I*_0_ is the transistor dark current which are constants of the n-type MOSFETs used in the circuit [42]. The non-linear term comes from *f* (*I*_*mem*_) which has been experimentally shown to be an exponential function of *I*_*mem*_ [43]. *I*_*τ*_ and *I*_*th*_ are bias currents generated from respective supply voltages using on-board bias generator circuits [44, 45].

The synapses of the DYNAP-SE2 are modeled after the DPI circuit [46] and can be represented in a simplified form as

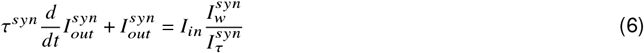

where 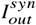 is the output current of the synapse, the time constant is 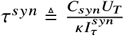, where 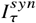 is a bias current, and 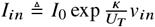 is the input current to the synapse.

### 5.2 AFE and encoding setup

The dataset chosen in this work consisted of intracranially recorded EEG (iEEG) signals. In the SWEC dataset, the iEEG signals are made available after undergoing 16-bit analog-to-digital conversion, and being band-pass filtered between 0.5 and 150 Hz, were recorded digitally at a sampling rate of 512 Hz. To mimic a real-world scenario, we utilized a DAC stage (acting as an EEG electrode placeholder) to convert the digitally recorded waveforms of the dataset into analog signals. This DAC stage consists of an 18-bit DAC (AD5781, Analog Devices, Wilmington, MA) and a custom PCB (designed with support from the Microelectronics Design Center of ETH Zurich), allowing regeneration of the digital data into corresponding analog waveforms (Fig. 6a). The DAC is controlled by a microcontroller (Teensy 4.1 from PJRC, Sherwood, OR) which transmits a digital signal stored in an SD card to the DAC via an SPI bus. The input signal is generated with a ∼ 100 mV peak-to-peak voltage and with a ∼ 30 mV offset for all DAC conversions. The EEG signal amplitude is scaled to such values considering the DAC PCB and AFE characteristics. A sample from the oscilloscope is presented in Fig. 6b, where the analog signal (trace 4; purple) can be seen. Note, the SWEC data has been converted to analog signals as downloaded without any pre-processing, in this study, by the DAC system described above.

The AFE signal conditioning stage consisted of a high-fidelity low-noise signal amplifier and a filter bank as depicted in Fig. 6c. We employed a 20 dB default amplification with no filtering. Note that the input signal (from the database) is pre-filtered to be within the 0.5 to 150 Hz frequency band at the source. This amplified signal is then passed on to the ADM. Although both the AFE and SNN cores are located within the same die, we have extracted the output events from the ADM sequentially, i.e., channel by channel, (for limited I/O of the AFE on the die as compared to the number of input EEG channels) before sending these simultaneously from off-chip to the SNN cores (Fig. 1f and Fig. 4b).

### 5.3 Delta modulated EEG events

Asynchronous Delta Modulation [47, 48] is a delta modulation approach where the sampling interval is adapted based on the characteristics of the coded signal. In the current context, it represents a level shifter circuit that outputs an event based on the difference in the amplitude of a signal between two sampling points. If two sample points (which might not be consecutively placed, representing asynchronous sampling obeying causality) of a signal show a change in amplitude that is more than a preset delta-threshold, the encoder records an event. An *‘UP’* or *‘DOWN’* tag is attached based on the polarity of the change, i.e. if the signal has increased or decreased at the level crossing. Further details on the circuit schematics of the AFE and the ADM can be found in Sharifshazileh et al. [29]. Due to the initial stabilization of analog components within the AFE circuit, we obtained 320 second snippets of each channel for a given seizure following a discarded two-minute pre-seizure segment.

### 5.4 Analog neurons with digital routing

Figure 7 shows the block diagram of DYNAP-SE2 comprising four neuron cores and the AFE. Each core contains 256 Adaptive Exponential Leaky-Integrate-and-Fire (AdEx I&F) neurons, totaling 1024 (Fig. 7a). All system parameters, such as time-constants, and gains of the synaptic compartments and threshold, and refractory periods of soma are set by an on-chip bias-generator [44, 45] and are shared within a single core. In other words, only four sets of parameters can be assigned within the entire chip. In this work, we have utilised three neural cores hosting R-E/I, FF-I and NLNG neurons. These parameters are configured as currents that map to a corresponding value of the parameter. This current are programmed through a *coarse* (*C*_*p*_ ∈ {0..5}) and a *fine* (*F*_*p*_ ∈ {0..255}) value, which follows the relation *I* = *I*_*c*_{*C*_*p*_}(*F*_*p*_/256) where *I*_*c*_ is estimated to be related with the *C*_*p*_ as described by Table 3 [49].

**Table 3.** Estimated current values corresponding to *coarse* bias values from Biasgen circuit simulations.

| $C_p$ | 0 | 1 | 2 | 3 | 4 | 5 |
| --- | --- | --- | --- | --- | --- | --- |
| $I_p$ (nA) | 0.07 | 0.55 | 4.45 | 35 | 280 | 2250 |

**Figure 7.**
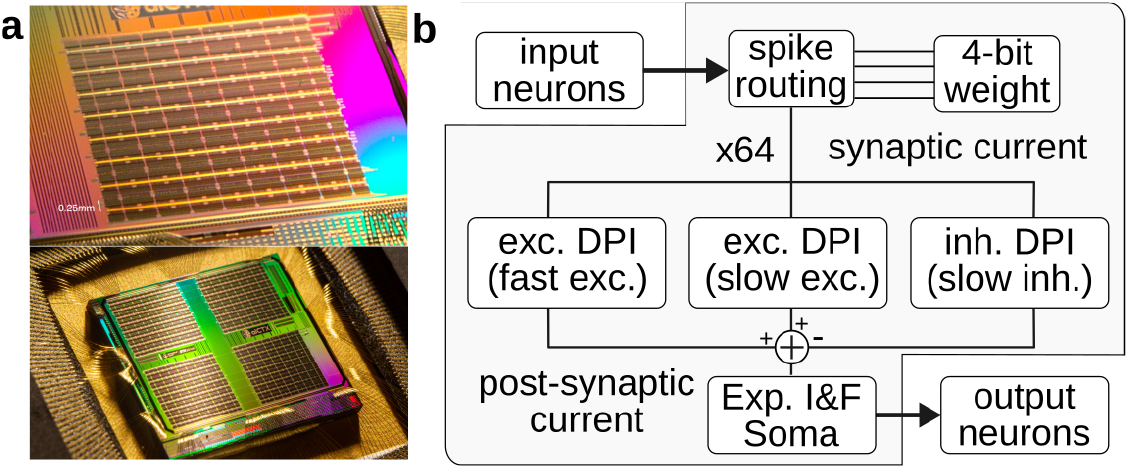
Analog neuron circuits in the DYNAP-SE2 Neuromorphic Processor; **a** Die photograph of the DYNAP-SE2 with (top) close-up shot of one of the neural cores; (bottom) and total four neural cores in a die (see Richter et al. [26] for details); **b** Neuronal block; Input events are translated to synaptic currents. The synaptic currents are routed to one of four synapse types, each combined with Differential Pair Integrators, and forwarded to an Adaptive Exponential Leaky Integrate & Fire soma.

Within this work, all biases for parameters except for synaptic parameters mentioned in Table 4 unchanged and at default recommended (*C, F*) values. Each neuron has 64 synapses (fan-in) that can receive spikes from any neuron throughout the whole chip and can be connected to up to 1024 post-synaptic (fan-out) neurons. In the event of an incoming spike, up to four synaptic currents are generated, their magnitudes based on 4-bit weights, that are added and fed into one of four synapse types: fast excitatory (AMPA-like), slow excitatory (NMDA-like), slow inhibitory (GABA-B like) and fast inhibitory (GABA-A like) (that acts as a shunt) to the soma (Fig. 7b). The fast excitatory and inhibitory synapse types are not used in this work.

**Table 4.** Estimated bias generator currents on DYNAP-SE2 for setting synaptic compartment time constants and gains for (excitatory; NMDA like) and inhibitory; GABA-B like) for different utilised neural cores.

| Synaptic parameters<br>for neural core type |  | R - E/I<br>exc. / inh. | FF-I | NLNG |
| --- | --- | --- | --- | --- |
| Time-constant | $(C, F)$ | (1,20) / (1,30) | (2,20) | (1,12) |
| | $I_{\tau}$ (nA) | 0.628 / 0.667 | 4.528 | 0.597 |
| Gain | $(C, F)$ | (2,100) / (2,100) | (3,100) | (2,100) |
| | $I_{gain}$ (nA) | 4.84 / 4.84 | 35.39 | 4.84 |
| Weight | $(C, F)$ | (3,128) / ((3,128)+(3,128)) | (3,128) | (3,128) |
| | $I_{gain}$ (nA) | 35.5 / 2×35.5 | 35.5 | 35.5 |

Table 4 describes the estimated bias currents for synaptic parameters of different neural cores utilised in this work’s proposed SNN. Chicca et al. [21] describe the circuit relation of the bias parameters with the generated synaptic current within a neural core. Such values are chosen heuristically to achieve certain input/output behaviour (as discussed in Sec. 2.2) from a given neural core. Throughout the chip, a 20–30% coefficient of variation (CV) is observed with mean being the intended bias current value for all set bias parameters as a result of the subthreshold MOSFET design and the 180 nm manufacturing process. The values mentioned in the parametric table (Table 4) are an estimation and more insights into the hardware effects on parametric values can be found in Zendrikov et al. [39].

For each measurement, the number of equivalently weighted synapses, referred to as the “weights” in the SNN, are set to ensure that the membrane potential reaches a value close to the threshold. In the case of the SNN in this work, the number of equivalent connections of the translation layer is set to ten throughout all translation layer synapses and one or two in the NLNG layer. Such heterogeneous neuromorphic processing systems with mixed-signal analog/digital electronic circuits are extremely noisy and imprecise owing to their sub-threshold operation with very small currents. Section 5.1 mentions the expressions relating the bias currents to their intended physical values for both synapses and neurons in a neural core of DYNAP-SE2. Circuit diagrams of the DPI circuits and a description of additional features for DYNAP-SE2 can be found in Richter et al. [26].

### 5.5 Power Consumption for SNN

Power consumption for analog silicon neurons with digital routing in the DYNAP family of neuromorphic processors can be estimated to be as follows (See Risi et al. [50], and Moradi et al. [51] for details):

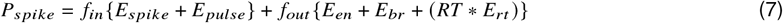

where,

*f*_*in*,*out*_ = Incoming and outgoing firing rate, respectively

*E*_*s pike*_ = 883 *pJ*, energy to generate one spike

*E* _*pulse*_ = 324 *pJ*, energy of the pulse extender circuit

*E*_*en*_ = 883 *pJ*, energy to encode one spike and append destination

*E*_*br*_ = 6840 *pJ*, energy to broadcast event to same core

*E*_*rt*_ = 360 *pJ*, energy to route event to different core

*RT* = 1, if the spike is sent to a different core, zero otherwise

This power estimation can be rewritten for the spike trains of different neuronal types (ADM, EE, RI, FF-I, NLNG) per iEEG channels, utilized in our hardware-implemented SNN. Note that, owing to the shared parameter within a neural core, R-E/I neurons, FF-I neurons, and NLNG neurons are implemented in different cores of the DYNAP-SE2 neuromorphic processor, respectively.

For ADM/AFE spikes,

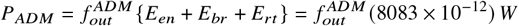

For neurons in the translation layer, for incoming spikes to both R-E and FF-I neurons

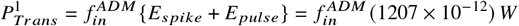

from R-E neurons, outgoing to NLNG neurons (in different core) and R-I neurons (in the same core)

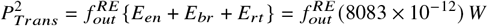

from FF-I neurons, outgoing to R-E neurons (in different core)

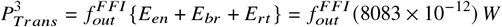

for R-I neurons, incoming spikes from RE neurons and outgoing to R-E neurons (in same core)

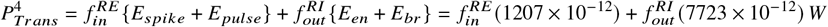

For neurons in the NLNG layer, for incoming from the R-E neurons and outgoing to the readout

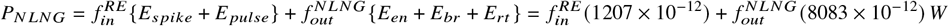

For our seizure encoding task, we divide the spike trains of each iEEG channel into four different windows, namely pre-seizure, first half (FH), and second half (SH) of the seizure and post-seizure in time, and calculate the spiking rate. From the calculated spike rate, the total power consumption and the average power consumption per channel are computed at the standard 1.8 V supply voltage for each seizure for each patient as presented in Table 5. We observe an average power consumption of 150 *μW* and 2.8 *μW*/per channel for encoding of the seizures by the SNN.

**Table 5.**
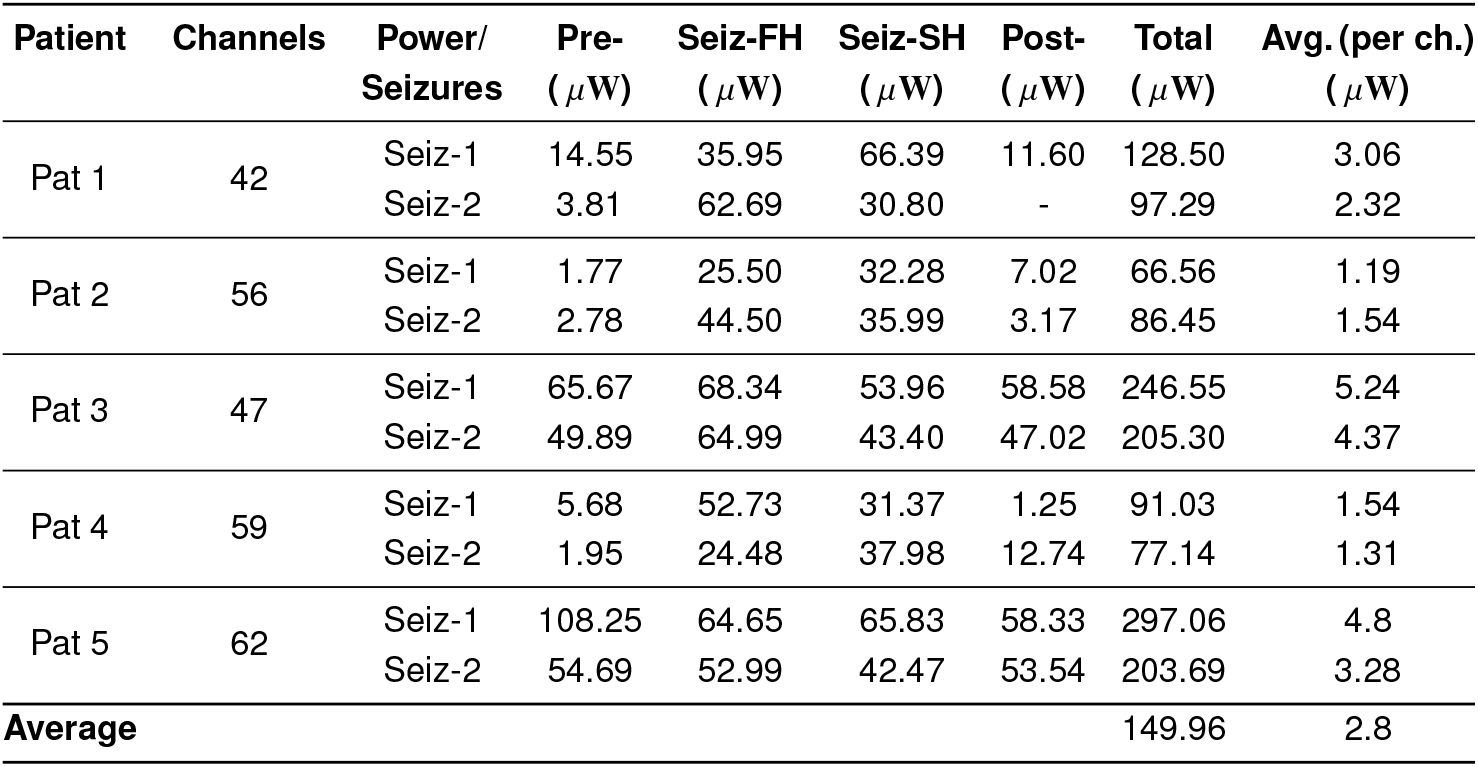
Power consumption for the SNN across patients.

| Patient | Channels | Power/<br>Seizures | Pre-<br>(μW) | Seiz-FH<br>(μW) | Seiz-SH<br>(μW) | Post-<br>(μW) | Total<br>(μW) | Avg. (per ch.)<br>(μW) |
| --- | --- | --- | --- | --- | --- | --- | --- | --- |
| Pat 1 | 42 | Seiz-1 | 14.55 | 35.95 | 66.39 | 11.60 | 128.50 | 3.06 |
|  |  | Seiz-2 | 3.81 | 62.69 | 30.80 | - | 97.29 | 2.32 |
| Pat 2 | 56 | Seiz-1 | 1.77 | 25.50 | 32.28 | 7.02 | 66.56 | 1.19 |
|  |  | Seiz-2 | 2.78 | 44.50 | 35.99 | 3.17 | 86.45 | 1.54 |
| Pat 3 | 47 | Seiz-1 | 65.67 | 68.34 | 53.96 | 58.58 | 246.55 | 5.24 |
|  |  | Seiz-2 | 49.89 | 64.99 | 43.40 | 47.02 | 205.30 | 4.37 |
| Pat 4 | 59 | Seiz-1 | 5.68 | 52.73 | 31.37 | 1.25 | 91.03 | 1.54 |
|  |  | Seiz-2 | 1.95 | 24.48 | 37.98 | 12.74 | 77.14 | 1.31 |
| Pat 5 | 62 | Seiz-1 | 108.25 | 64.65 | 65.83 | 58.33 | 297.06 | 4.8 |
|  |  | Seiz-2 | 54.69 | 52.99 | 42.47 | 53.54 | 203.69 | 3.28 |
| Average |  |  |  |  |  |  | 149.96 | 2.8 |

### 5.6 Spike Train Correlations

To quantitatively determine the correlated state of ADM event steams and NLNG neurons across a given time window, namely, synchronization (as interpreted in Sec. 2.3), we have calculated the Manhattan (*L*_1_) norm of the inner product between kernelized spike trains from NLNG neuron pairs and ADM event-stream-pairs..

#### 5.6.1 Kernelizing spike-trains to time-continuous functions

The time-binned binary representation of event timestamps can lead to loss of information depending on the choice of time bin widths [37]. Here, we used a kernelization approach to transform discrete time-stamps of events into a time-continuous signal [37, 52].

For a neuron in NLNG layer or a event steam from ADM, the spike train can be denoted as 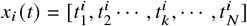, with 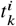 being time-stamps of the event-steam *i* with a total of *N* event time-stamps. Such discrete series can be alternatively represented as summation of Dirac Delta functions: 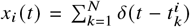, where *i* = [1, 2, · · ·, *M*] with *M* being the total number of channels under consideration (See Fig. 3). Thereafter, this spike trains can be transformed into a time-continuous signal *f*_*i*_ (*t*) though a convolution *h*, described by:

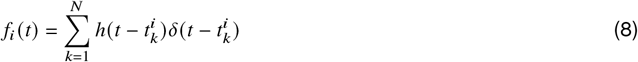

Here, *i* represents the spike train of neuron or event-stream from ADM *i* with *i* = [1, 2, · · ·, *M*] .. *h* is the kernel, represented by a causal exponential decay

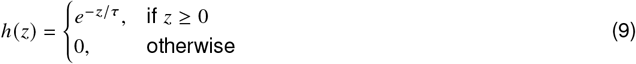

with *τ* being the time constant assigned to the kernel *h*. Here, *τ* is set to 1 ms to include the long and short-term effects of spikes. Further details can be found in Park. et al [53].

#### 5.6.2 Manhattan norm and synchronization matrices

Next, for a set of *M* spike train functions, with *M* representing the number of total spike trains (or channels):

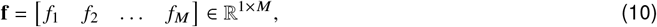

where each *f*_*i*_ is associated with the kernelized spike-train of neuron *i*, defined by Eq. 8. From the set **f**, we can define a cross-correlation matrix **C** = **ff**^*T*^, ∈ ℝ^*M*×*M* 1^, where each element is given by

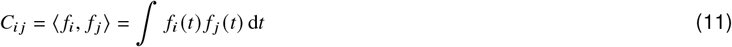

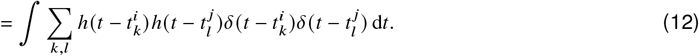

Thus, the resulting matrix **C** ^2^ **is given by**

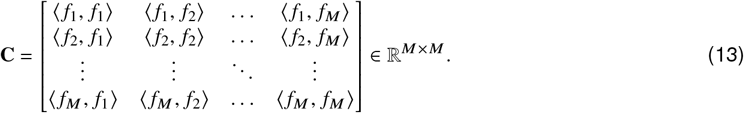

**Now, for a given time window *T***, **We can define C** as **N** ∈ ℝ ^*M*×*M*^, such that its elements are given by

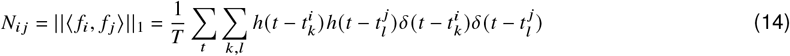

We further set *N*_*i*,*i*_ = 0 throughout the analysis. This Manhattan (*L*_1_) norm of the inner product, averaged over time *T*, serves as a measure of synchronization (See Sec. 2.3). The considered time window T is chosen to be half of the seizure window-length in Fig. 4, full seizure window-length in Fig. 5 and a non-overlapping rolling time window of 1 sec in Sec. 2.4.

#### 5.6.3 Support Vector Machines (SVM)

For the seizure detection task as a post-processing step, we used standard Support Vector Machines (SVM), with linear and Radial Basis Function (RBF) kernels to classify seizure and non-seizure events on a per-patient basis. The linear SVM (LSVM) attempts to find the optimal hyperplane that best separates the two classes (seizure vs. non-seizure, labeled as 1 and 0 respectively) in the feature space. Specifically, given a set of data points (**x**_*i*_, *y*_*i*_), where **x**_*i*_ ∈ ℝ^*k*^ represents the *k* features (ADM or NLNG firing rates) of the i-th window, and *y*_*i*_ ∈ 0, 1 represents whether the window is a non-seizure or seizure event, the SVM algorithm searches for a separating hyperplane of the form **w** · **x**_*i*_ + *b* = 0. This hyperplane maximizes the margin between the two classes, where the margin is the distance between parallel hyperplanes **w** · **x**_*i*_ + *b* ≥ 1 for seizure windows and **w**· **x**_*i*_ + *b* ≤ 0 for non-seizure windows. For linear SVM, the regularization parameter *C* is included and controls the trade-off between maximizing the margin and allowing classification errors. A large value of *C* penalizes misclassifications more strictly, resulting in a smaller margin but fewer errors. In contrast, a smaller *C* allows for a larger margin with more flexibility. For the RBF kernel, an additional hyperparameter *γ* is included that defines the influence of each data point on the decision boundary. For the hyperparameter optimization and binary classification procedure we utilized Python’s scikit-learn and scikit-optimize (skopt) libraries.

#### 5.6.4 Performance Metrics

To evaluate the performance of the seizure detection models, we utilized the following standard classification metrics:

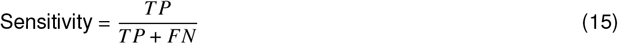

where true positives (TP) are seizure windows correctly classified as seizures, and false negatives (FN) are seizure windows incorrectly classified as non-seizures.

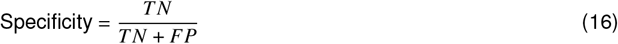

where true negatives (TN) are non-seizure windows correctly classified as non-seizures, and false positives (FP) are non-seizure windows incorrectly classified as seizures.

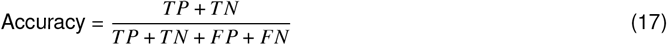

representing the overall proportion of correctly classified windows, both seizure and non-seizure.

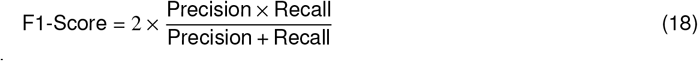

where Precision and Recall is defined as:

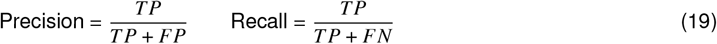

The F1-Score provides a balance between precision and recall, particularly useful for imbalanced datasets.

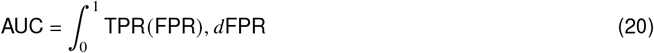

summarizing the trade-off between sensitivity (True Positive Rate) and 1-specificity (False Positive Rate) across different decision thresholds.

### 5.7 SWEC-ETHZ iEEG dataset

The intracranial EEG data used in this study (Table 6) were downloaded from a public epilepsy dataset [32], assembled at the Sleep-Wake-Epilepsy-Center (SWEC) of the University Department of Neurology at the Inselspital Bern and the Integrated Systems Laboratory of the ETH Zurich, available from http://ieeg-swez.ethz.ch/. Detailed information about the acquisition methods and patient metadata is contained in Refs. [54, 55]. The iEEG dataset is recorded during pre-surgical evaluation for epilepsy brain surgery. The intracranial EEG was recorded with strip, grid, and depth electrodes. The data was pre-filtered to the 0.5 to 150 Hz frequency band, sampled at 512 Hz, and further preprocessed to remove artifact-corrupted channels. Annotated seizure onsets and offsets were obtained from visual inspection of an EEG board-certified, experienced epileptologist. Readers are referred to the http://ieegswez.ethz.ch/ and associated publications for details about the pre-processing stages of the recorded EEG data. Each patient has recordings of multiple seizure events consisting of three minutes of pre-seizure segments (system state immediately before the seizure onset), the seizure segment (seizure event ranging from 10 s to 1002 s), and three minutes of post-seizure segment (i.e., system state after seizure event). All the information about the dataset was taken from the dataset website [32] and the associated publications [54, 55].

**Table 6.** Patient characteristics. TLE Temporal Lobe Epilepsy, PLE Parietal Lobe Epilepsy.

| Patient | Age (y) | SWEC label | No. of electrodes | Seizure durations<br>Sez-1/Sez-2 (s) | Epilepsy<br>type |
| --- | --- | --- | --- | --- | --- |
| 1 | 19 | ID2 | 42 | 96 / 260 | TLE |
| 2 | 27 | ID9 | 56 | 104 / 118 | TLE |
| 3 | 24 | ID1 | 47 | 125 / 70 | TLE |
| 4 | 31 | ID16 | 59 | 81 / 67 | TLE |
| 5 | 32 | ID4 | 62 | 160 / 127 | PLE |

Due to hardware-related constraints described previously in the AFE section, we have chosen a subset of the short-term dataset containing five patients, 2 seizures per patient. Furthermore, considering a constraint on the number of simultaneous input spike trains to the SNN on the event-based processors [26], we have selected patient data with less than 64 electrodes.

## 6 Acknowledgements

We thank Alfonso Blanco Fontao and Microelectronics Design Center, D-ITET, ETH Zürich for providing design and assembly support for the DAC board. We are also thankful to Adrian Whatley, Maryada Maryada, Chenxi Wu, and other members of the NCS group and the INI admin team for providing a conducive research environment that made this work possible.

## 6.2 Author contributions

This work was conceptualized by J.B., S.G., O.G., and G.I. This work was prepared by J.B., O.G., S.G., and G.I. who also wrote the original draft text of the manuscript and helped in the preparation of the figures. All authors reviewed and edited parts of the manuscript.

## 6.4 Data availability

The dataset analyzed during the current study is a publicly available dataset at http://ieeg-swez.ethz.ch/ [32]. The event-based dataset, containing output spike-trains from the AFE and the NLNG layer along with the DAC-generated values of the analog EEG signals are published in https://doi.org/10.5281/zenodo.10800793 [33].

## 6.5 Code availability

Data processing and analysis were conducted in Python 3.*x* and the code has been made available at Ref. [33] and published in https://doi.org/10.5281/zenodo.10800793.

## Footnotes

1 Note, inner product of real-valued vectors (or kernelized spike trains, here) represent zero lag correlation of the vectors.

2 We note ⟨ *f*_*i*_, *f* _*j*_ ⟩ ≥ 0 across all *i* and *j* and ⟨ *f*_*i*_, *f* _*j*_ ⟩ = ⟨ *f* _*j*_, *f*_*i*_ ⟩, making it a symmetric matrix.

