## Supplementary material for "Event Driven Neural Network on a Mixed Signal Neuromorphic Processor for EEG Based Epileptic Seizure Detection": Supplimentary Material

### This supplementary file includes:

Figs. S1-10 Output rasterplots for all neuron types (ADM/AFE,RE,RI,FFI and NLNG) from DYNAP-SE2 for all patients and seizures. The raster plots are color codded by instantaneous firing rates with 1 sec non-overlapping time windows.

Figs. S11-15 The synchronization matrices of for all patients and seizures. The top row depicts the ADM/AFE channels and the bottom row depicts the non-local non-global (NLNG) neurons. The columns from left to right show synchronization during the pre-Seizure, first and second half of the Seizure, and post-Seizure periods respectively.

Figs. S16-17 Synchronization matrices of non-local non-global (NLNG) neurons for two seizures of patient 2, based on three distinct input channel ordering methods. The first, second, and third rows represent synchronization based on input with the original dataset order, maximum correlation of the original data, and randomized order, respectively.

Fig. S18 Mean synchronization over all pairs of the non-local non-global NLNG neurons cumulatively summed over time for all patients.

Fig. S19 The synchronization across all non-local non-global layer (NLNG) neuron pair over time for all patients and seizures.

Fig. S19-20 Averaged performance metrics across patients using SVM with Linear and RBF kernels based on the firing rates and synchronization measure, respectively for ADM/AFE input and NLNG output.

Fig. S21 Accuracy across step and window sizes using SVM with Linear and RBF kernels for ADM/AFE input and NLNG output.

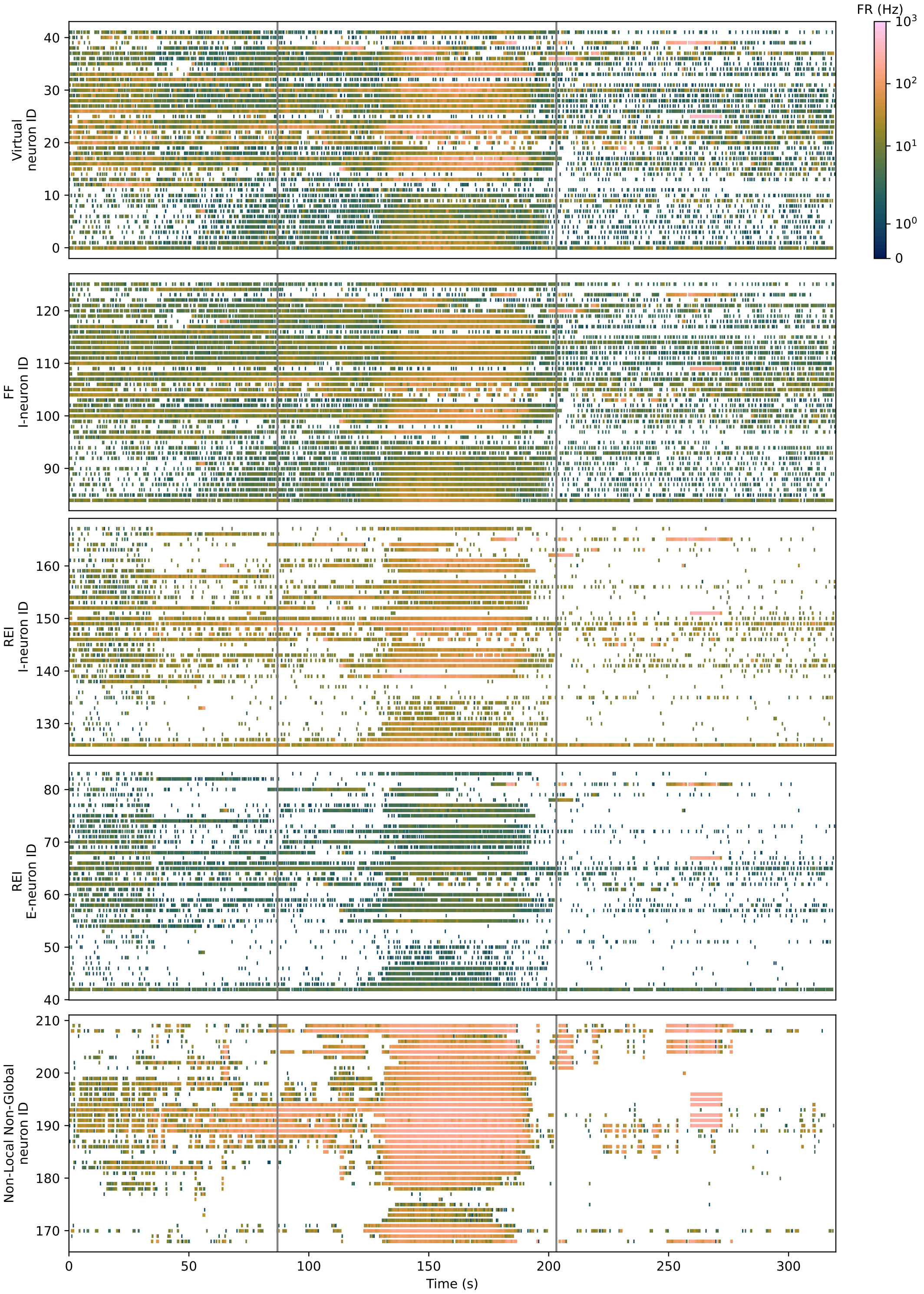


Supplementary Figure SF1: Patient 1 seizure 1 output rasterplot from DYNAP-SE2. Grey vertical lines indicate the start and end of the ictal period.

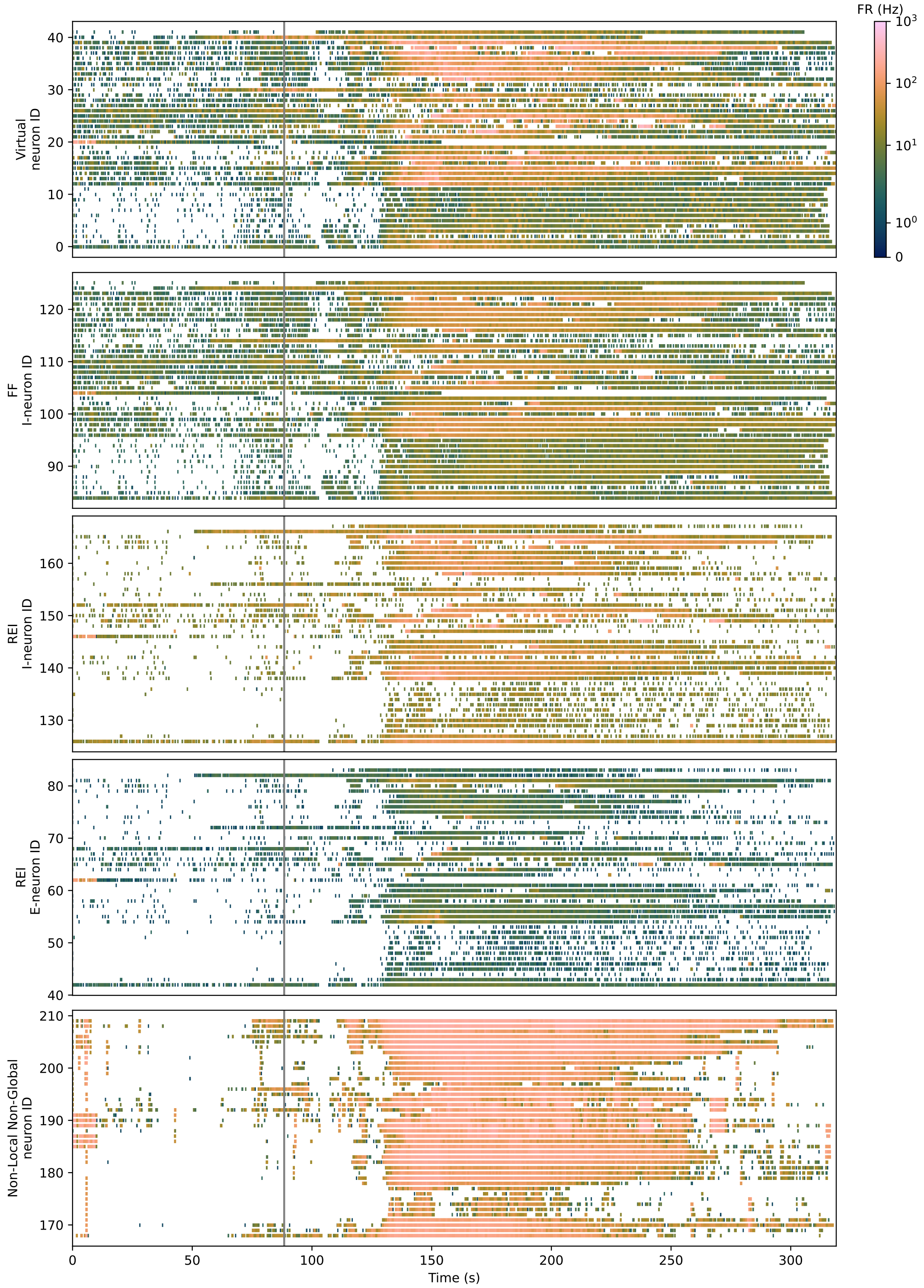


Supplementary Figure SF2: Patient 1 seizure 2 output rasterplot from DYNAP-SE2. Grey vertical line indicates the start of the ictal period.

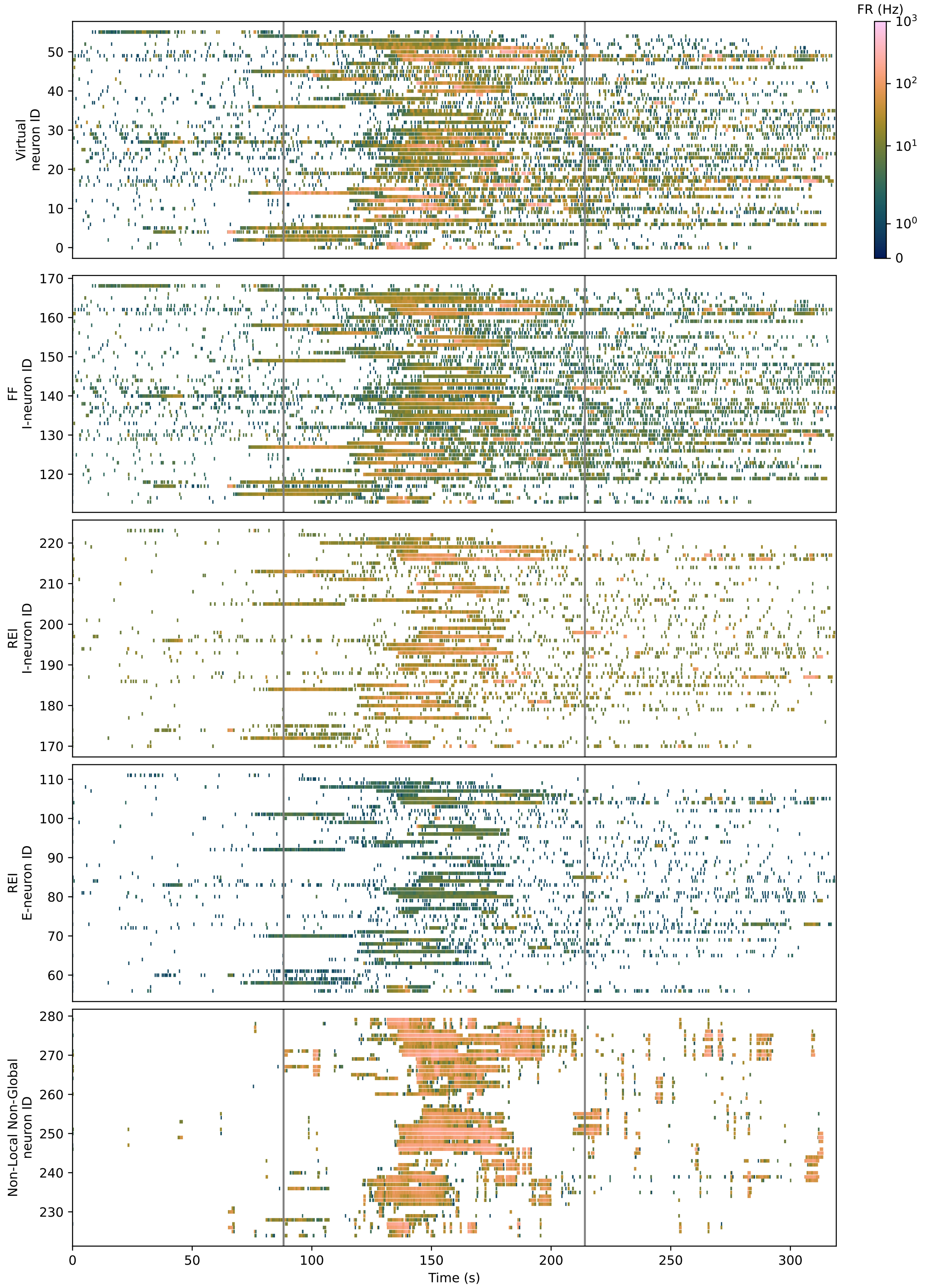


Supplementary Figure SF3: Patient 2 seizure 1 output rasterplot from DYNAP-SE2. Grey vertical lines indicate the start and end of the ictal period.

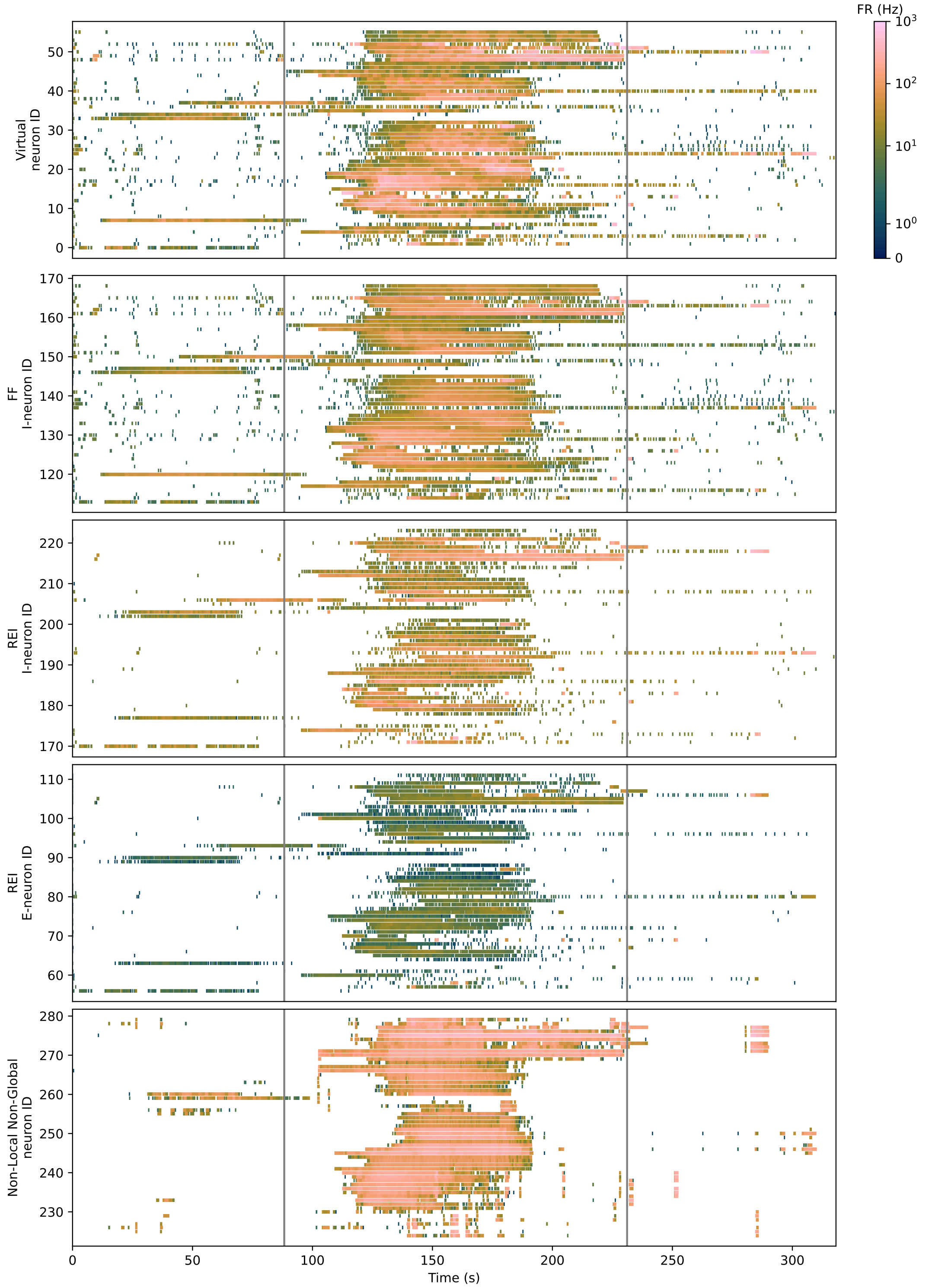


Supplementary Figure SF4: Patient 2 seizure 2 output rasterplot from DYNAP-SE2. Grey vertical lines indicate the start and end of the ictal period.

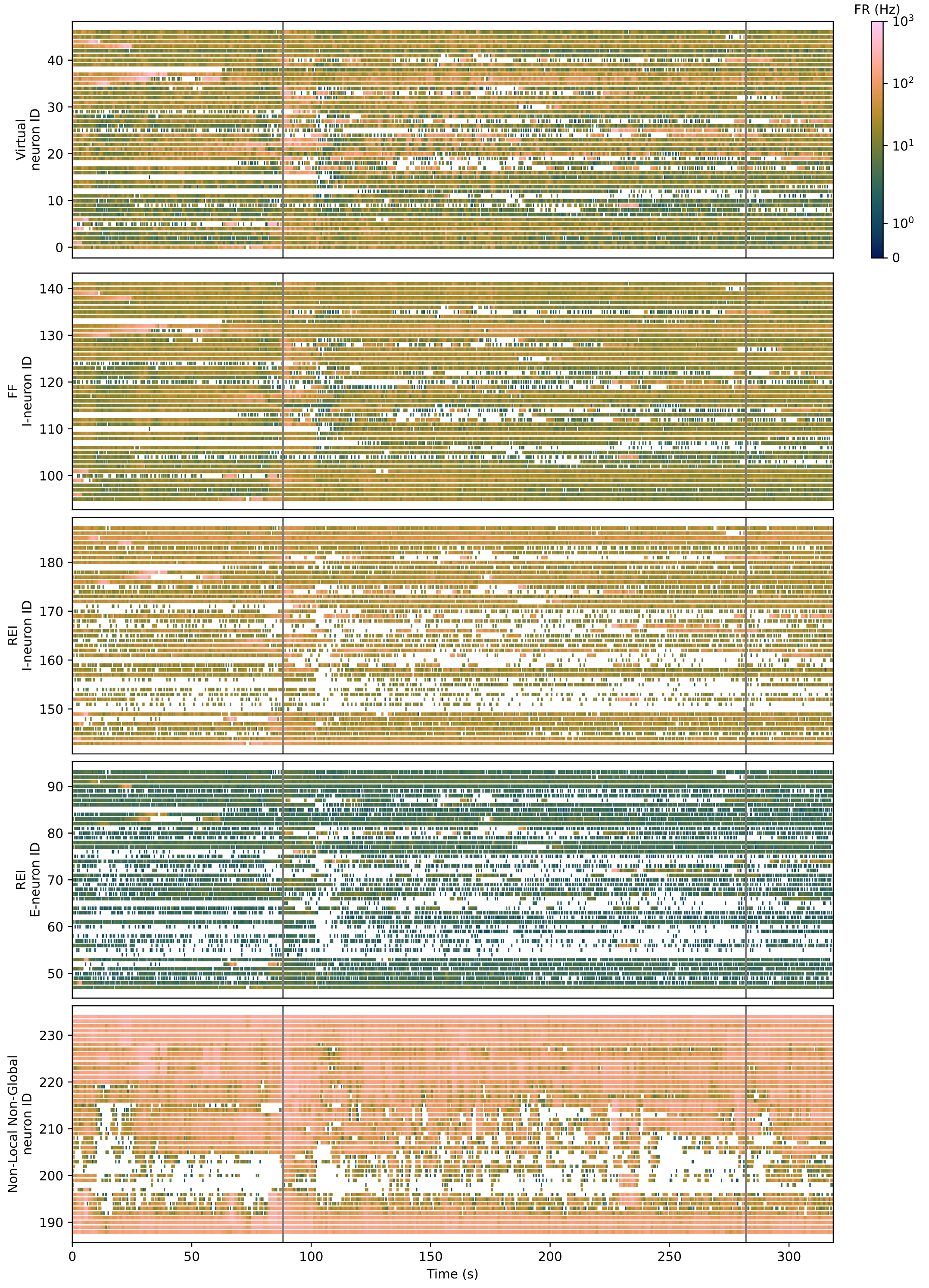


Supplementary Figure SF5: Patient 3 seizure 1 output rasterplot from DYNAP-SE2. Grey vertical lines indicate the start and end of the ictal period.

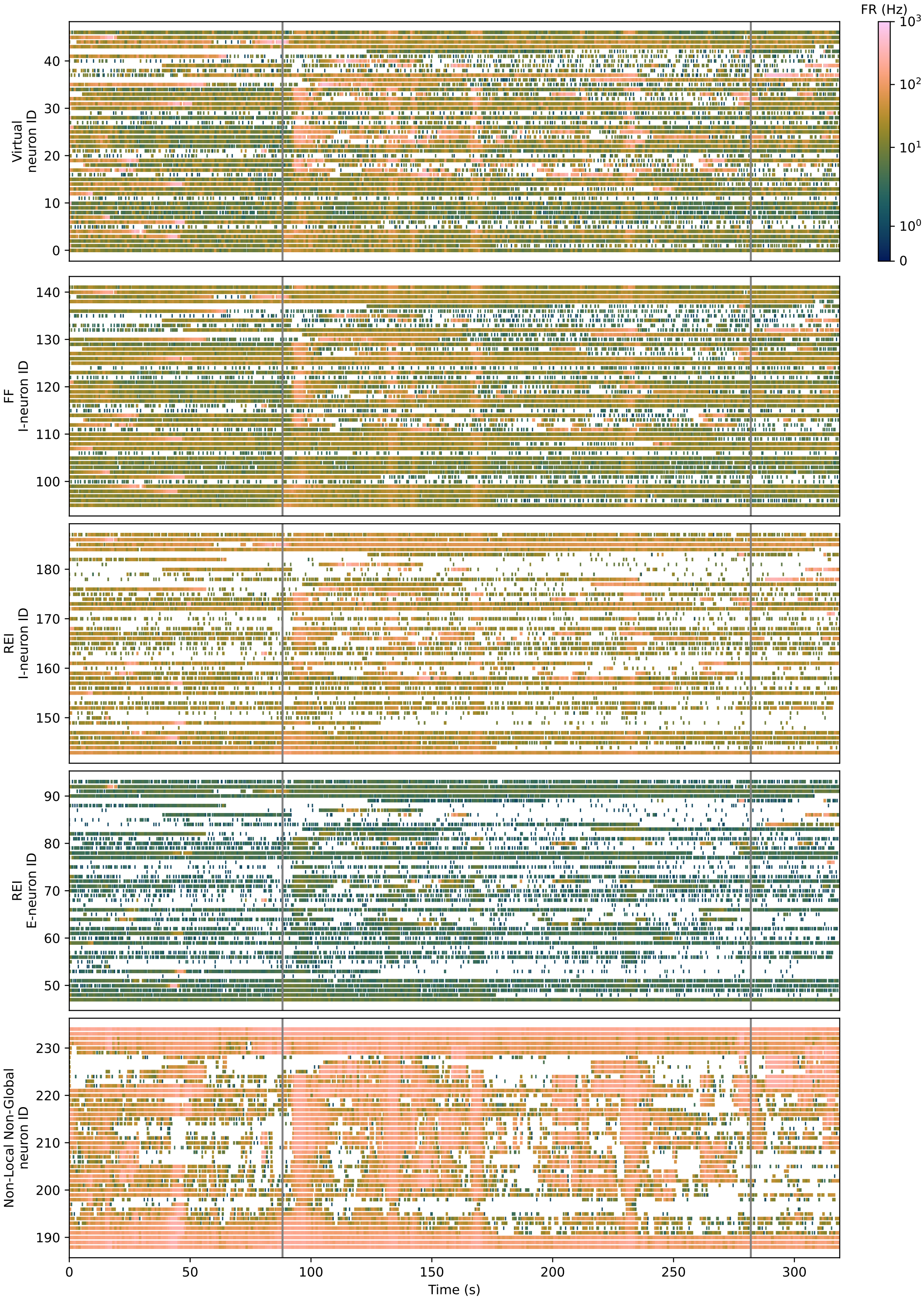


Supplementary Figure SF6: Patient 3 seizure 2 output rasterplot from DYNAP-SE2. Grey vertical lines indicate the start and end of the ictal period.

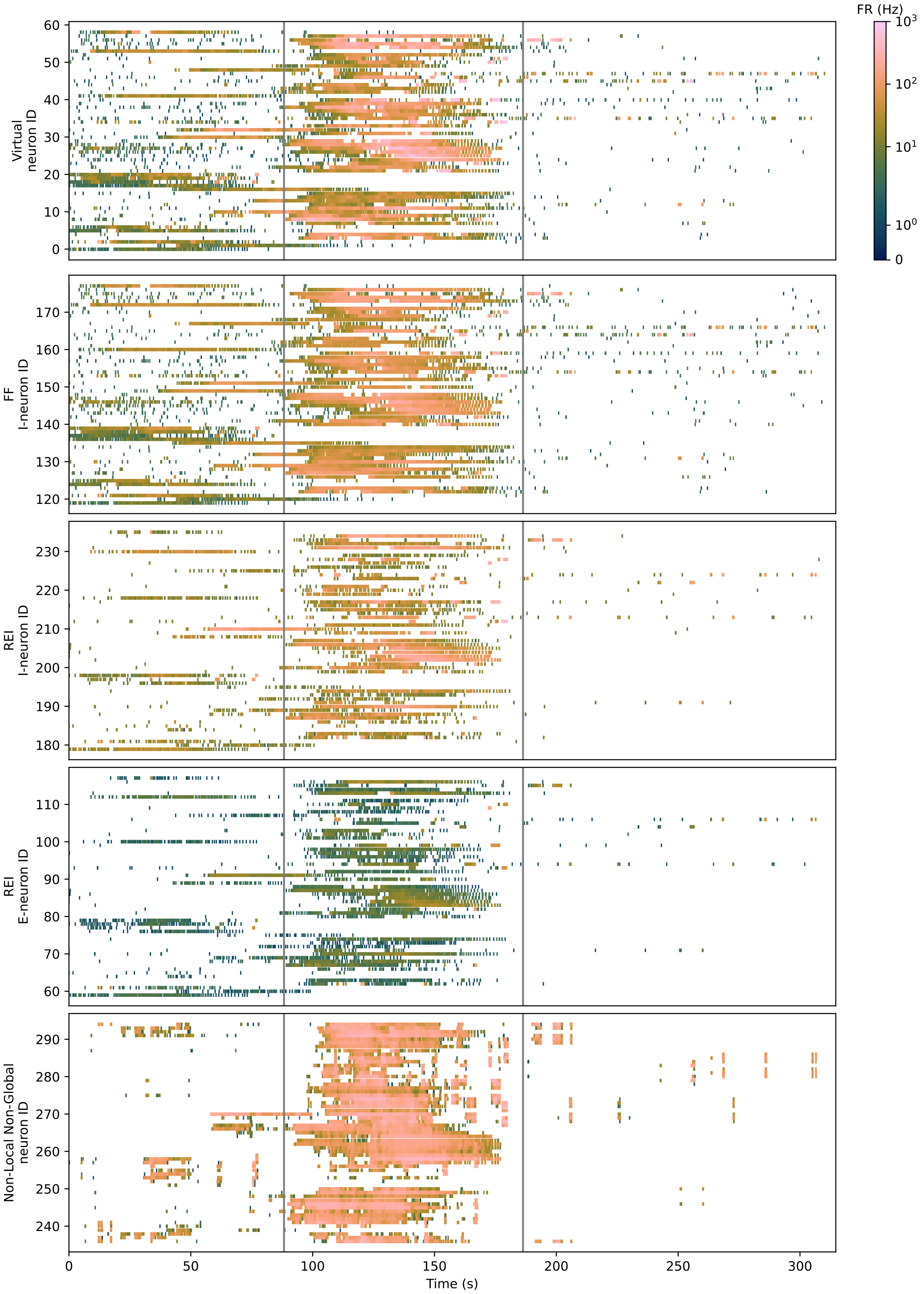


Supplementary Figure SF7: Patient 4 seizure 1 output rasterplot from DYNAP-SE2. Grey vertical lines indicate the start and end of the ictal period.

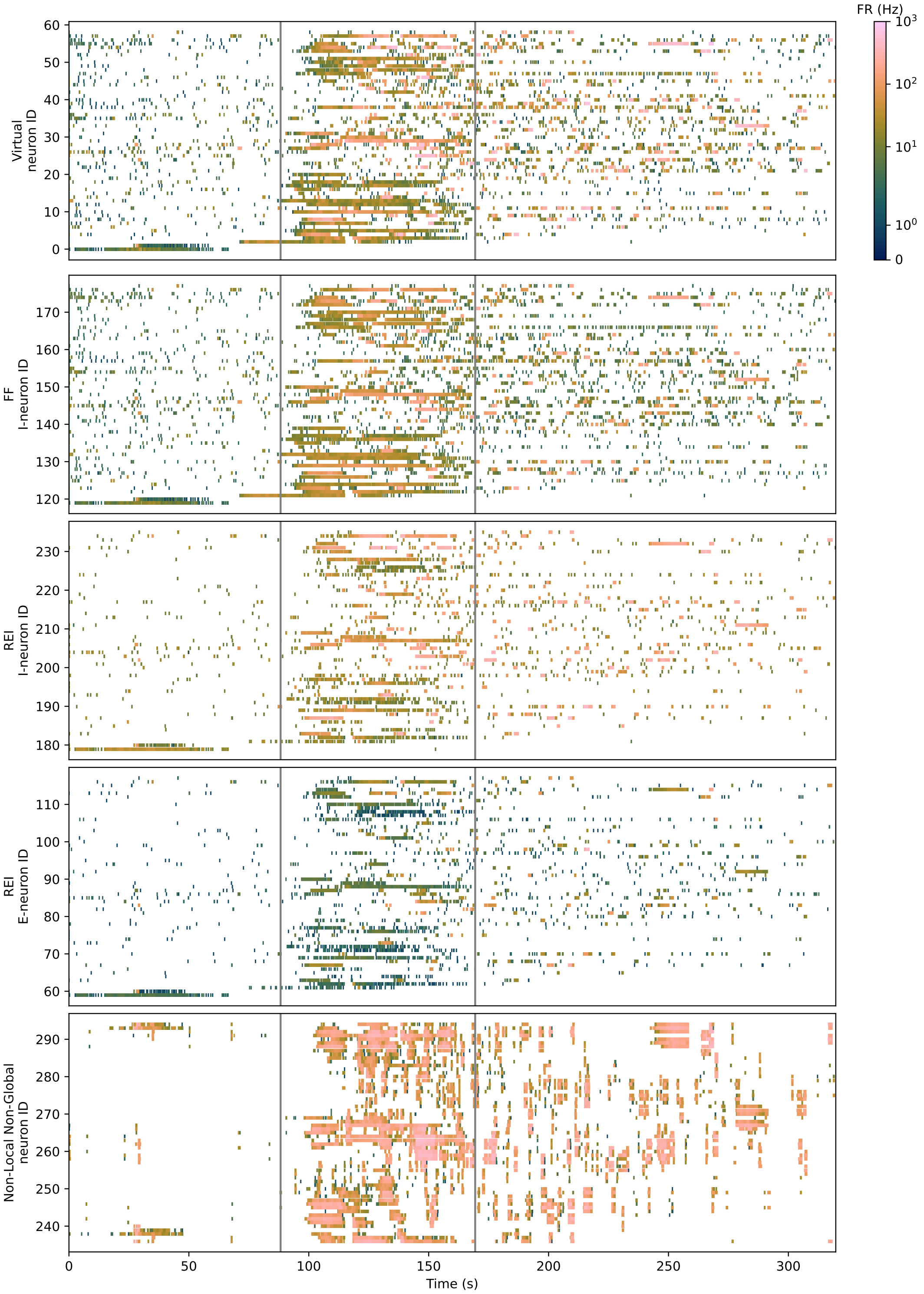


Supplementary Figure SF8: Patient 4 seizure 2 output rasterplot from DYNAP-SE2. Grey vertical lines indicate the start and end of the ictal period.

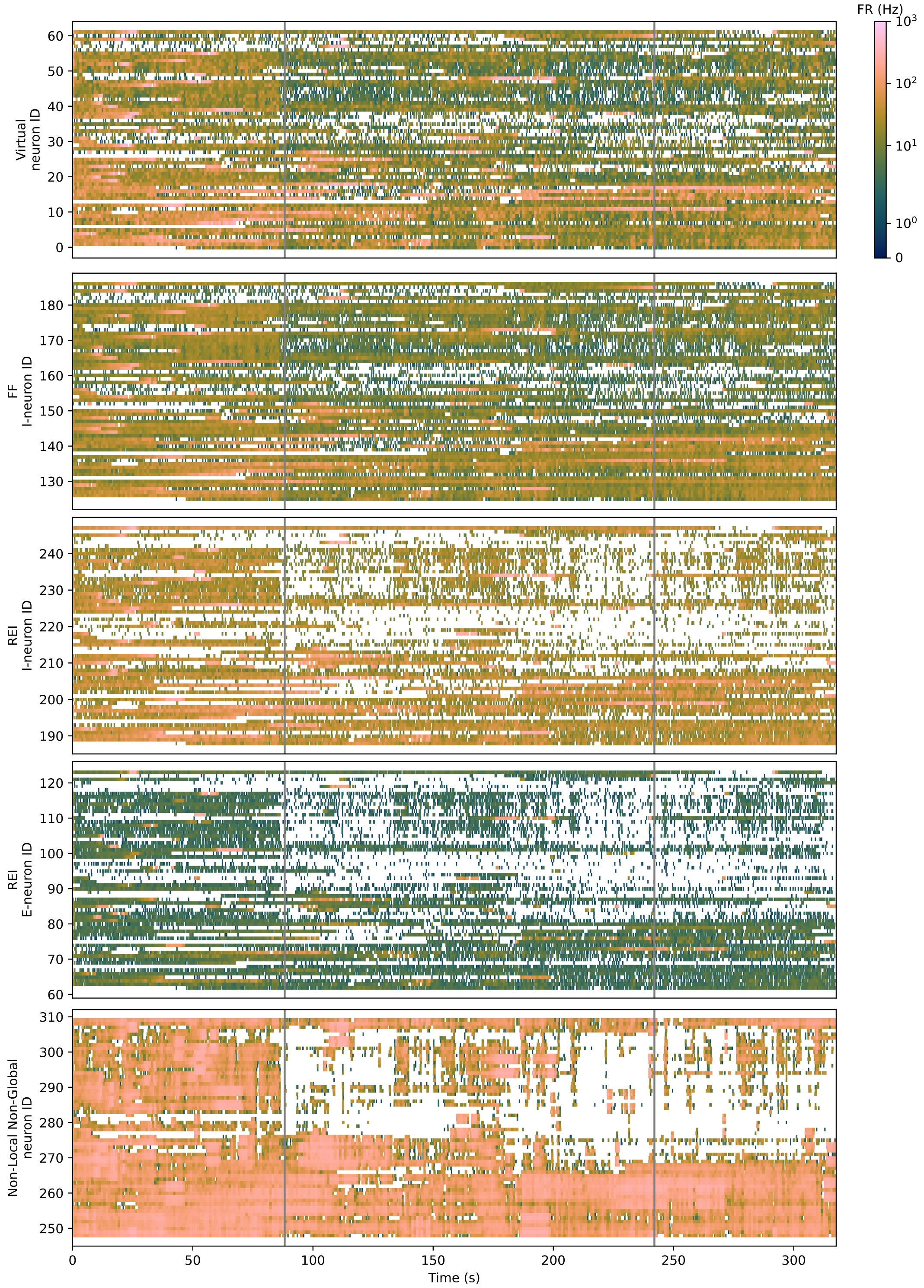


Supplementary Figure SF9: Patient 5 seizure 1 output rasterplot from DYNAP-SE2. Grey vertical lines indicate the start and end of the ictal period.

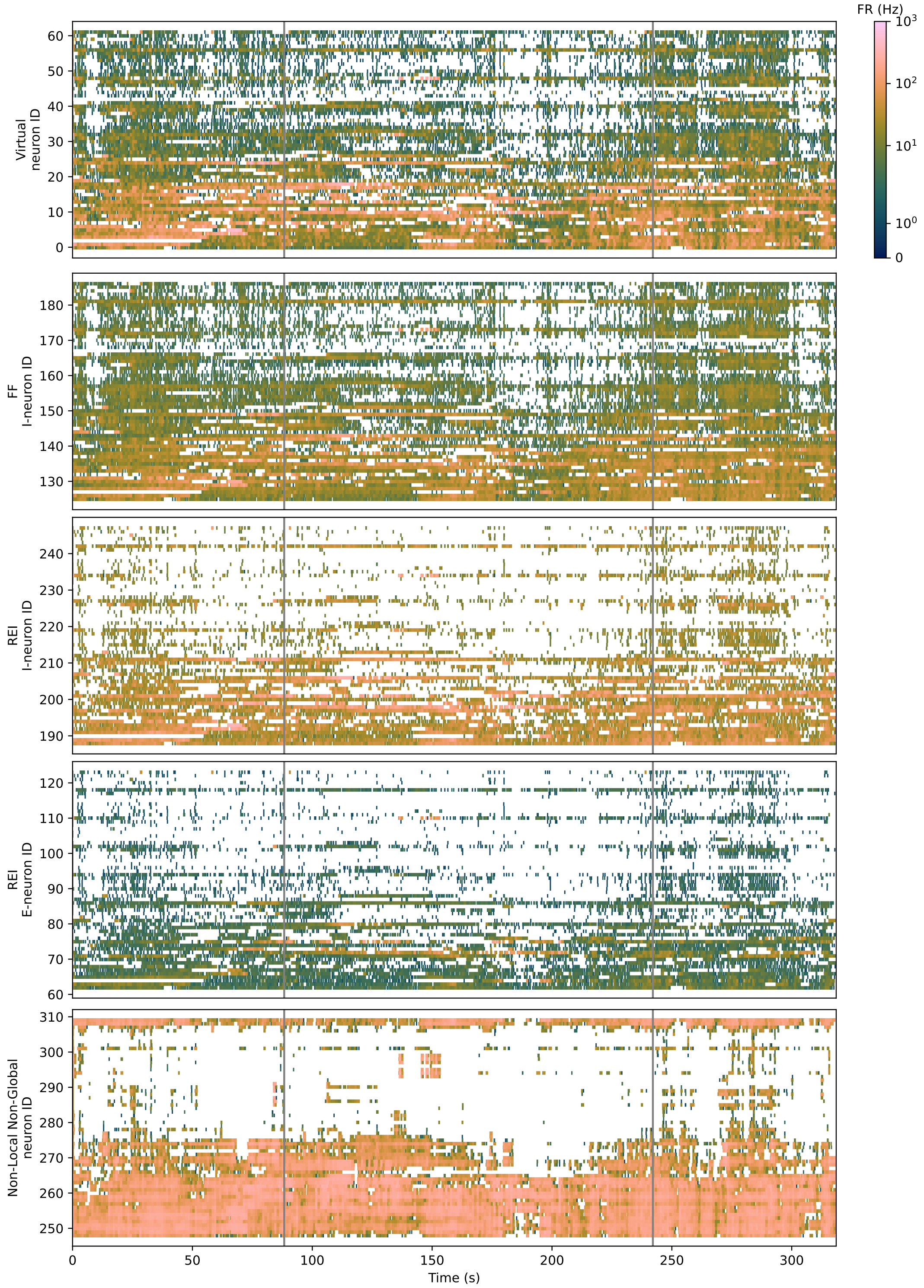


Supplementary Figure SF10: Patient 5 seizure 2 output rasterplot from DYNAP-SE2. Grey vertical lines indicate the start and end of the ictal period.

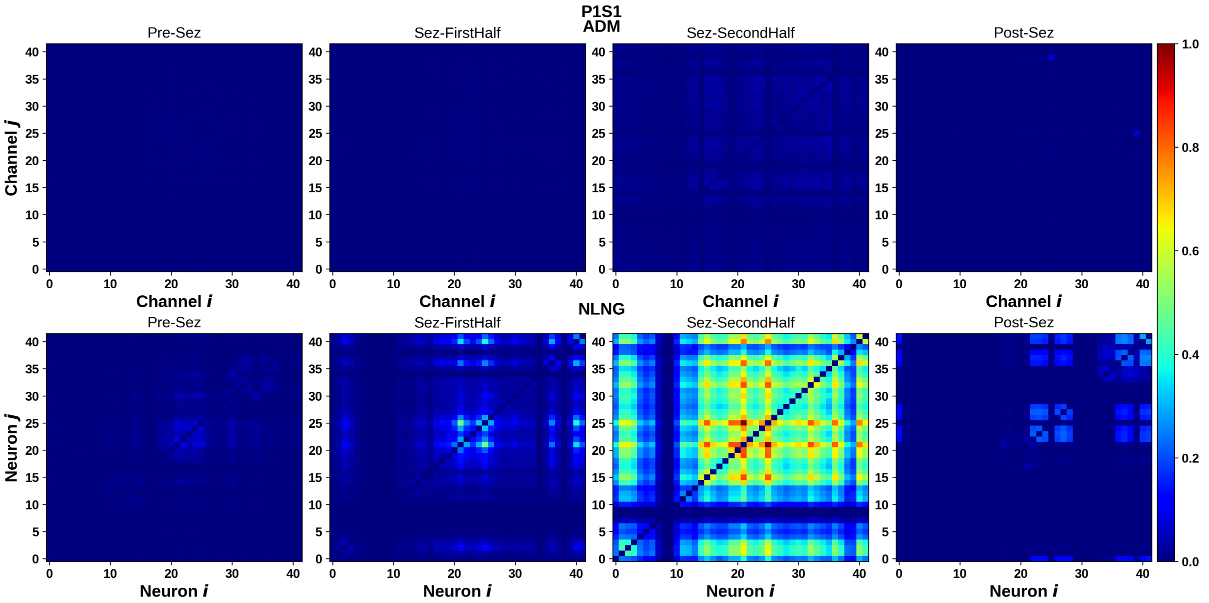


(a) Patient 1, Seizure 1

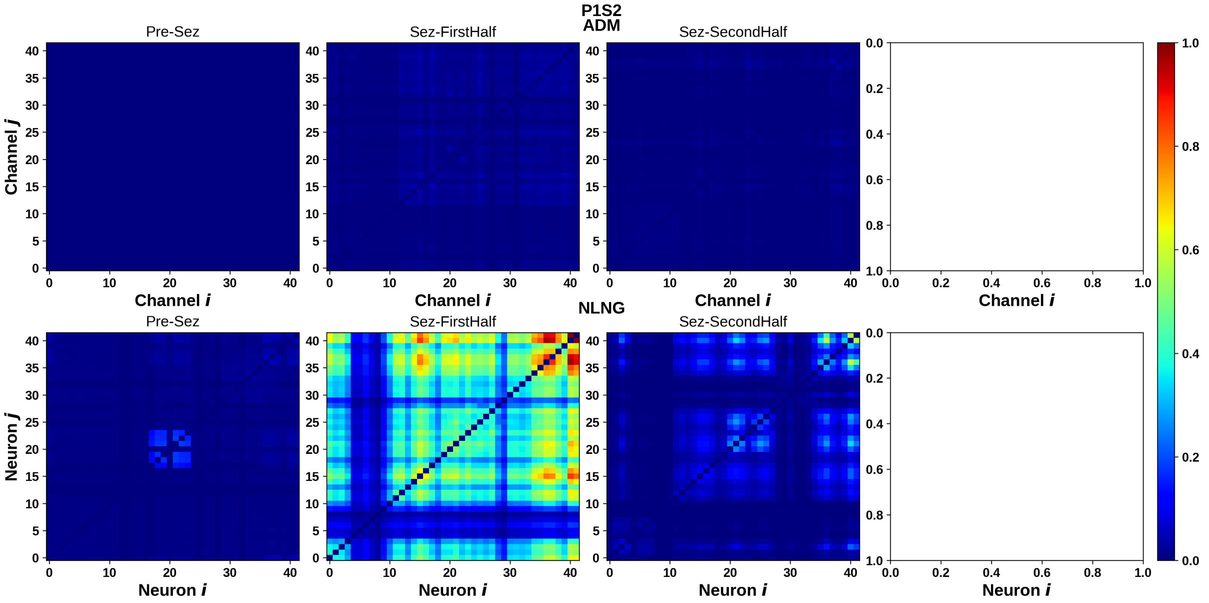


(b) Patient 1, Seizure 2

Supplementary Figure SF11: Synchronization matrices of Patient 1. The top row depicts the ADM/AFE channels and the bottom row depicts the NLNG neurons. The columns from left to right show synchronization during the pre-Seizure, first and second half of the Seizure, and post-Seizure periods respectively.

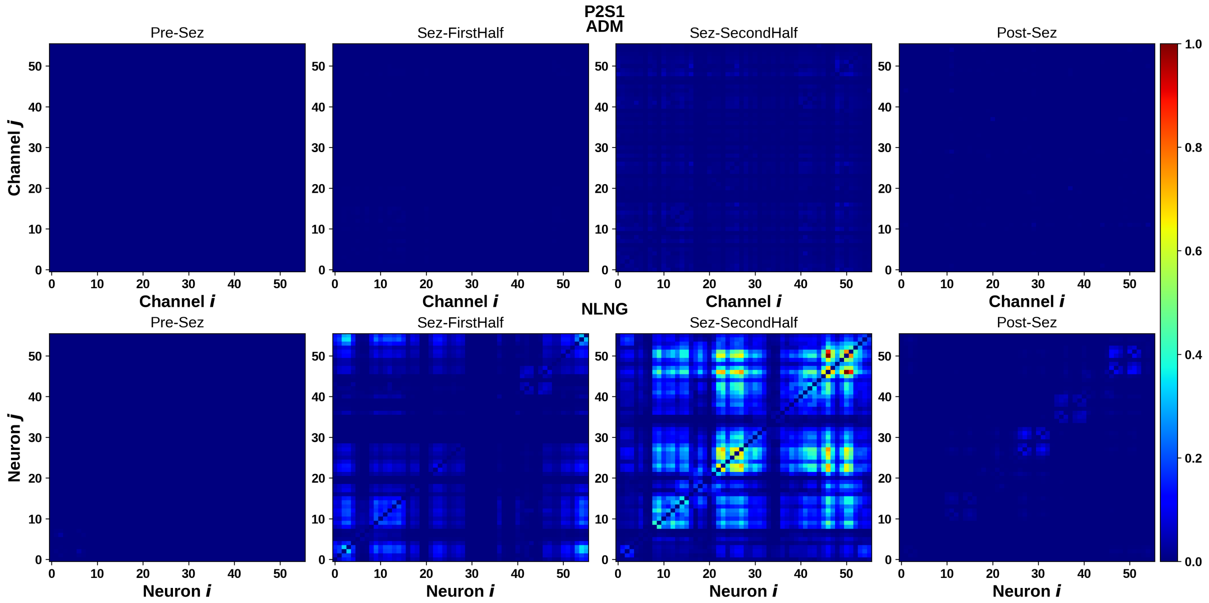


(a) Patient 2, Seizure 1

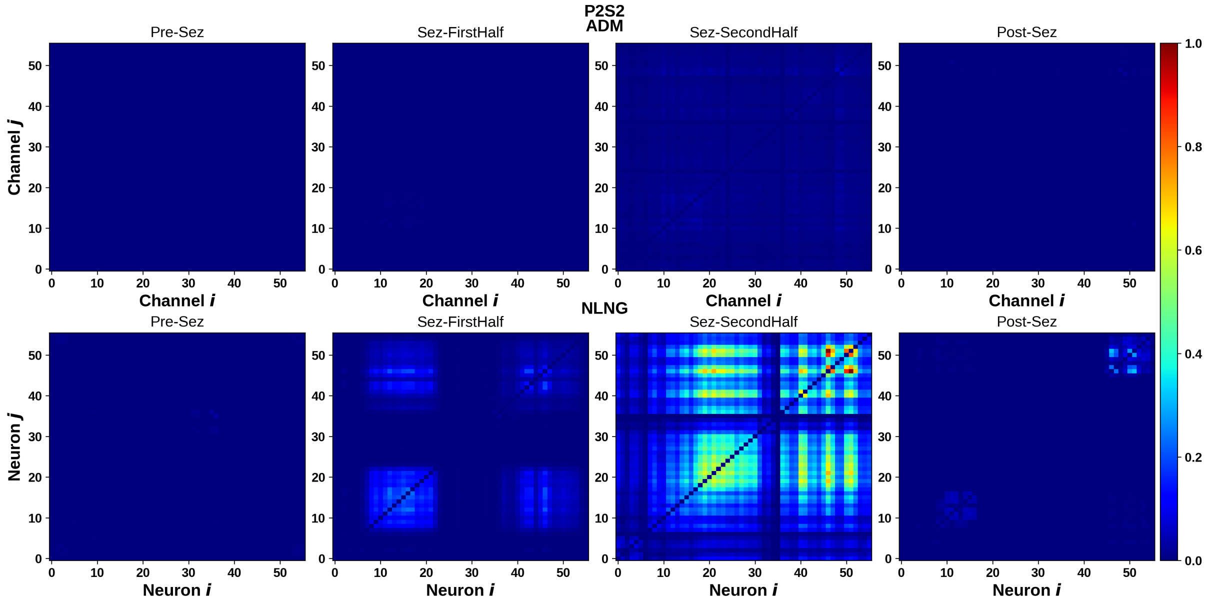


(b) Patient 2, Seizure 2

Supplementary Figure SF12: Synchronization matrices of Patient 2. The top row depicts the ADM/AFE channels and the bottom row depicts the NLNG neurons. The columns from left to right show synchronization during the pre-Seizure, first and second half of the Seizure, and post-Seizure periods respectively.

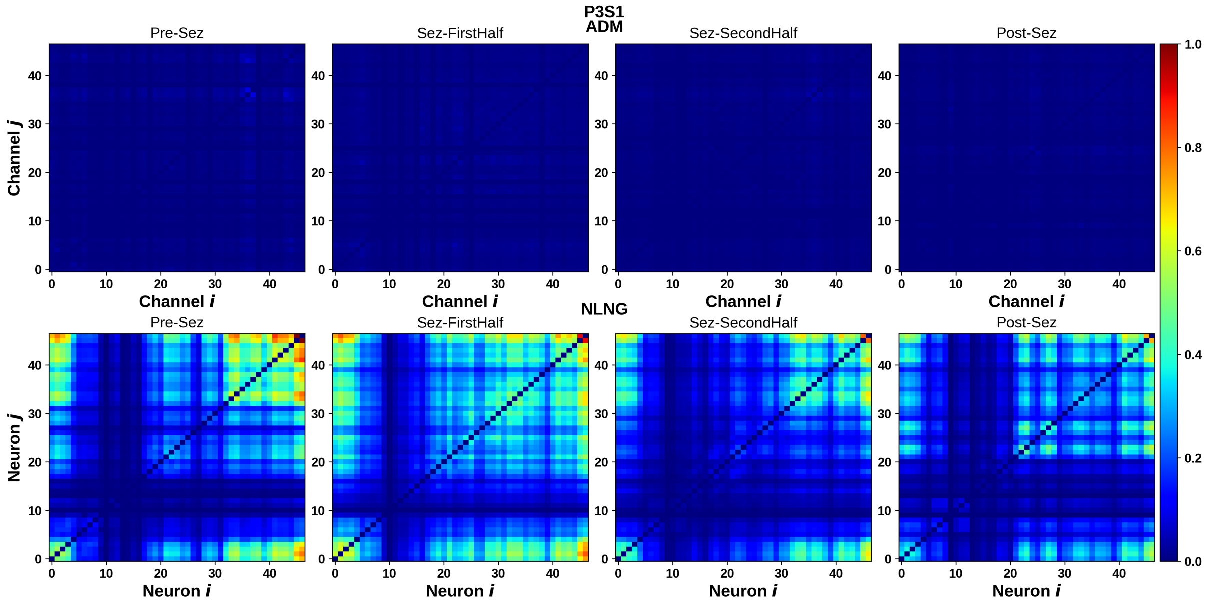


(a) Patient 3, Seizure 1

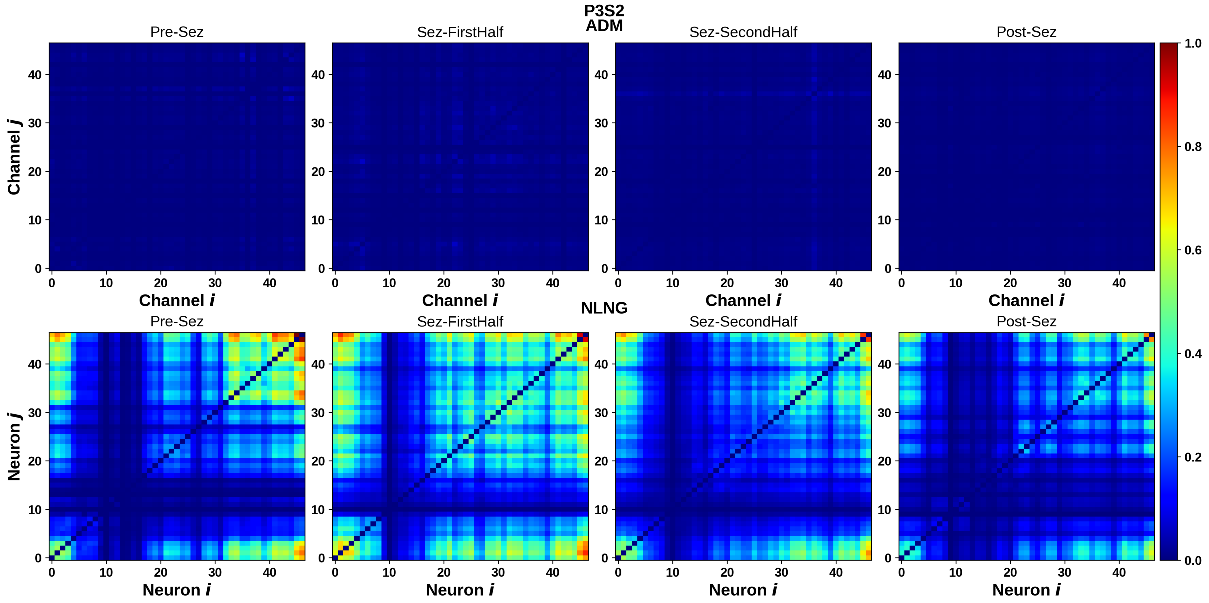


(b) Patient 3, Seizure 2

Supplementary Figure SF13: Synchronization matrices of Patient 3. The top row depicts the ADM/AFE channels and the bottom row depicts the NLNG neurons. The columns from left to right show synchronization during the pre-Seizure, first and second half of the Seizure, and post-Seizure periods respectively.

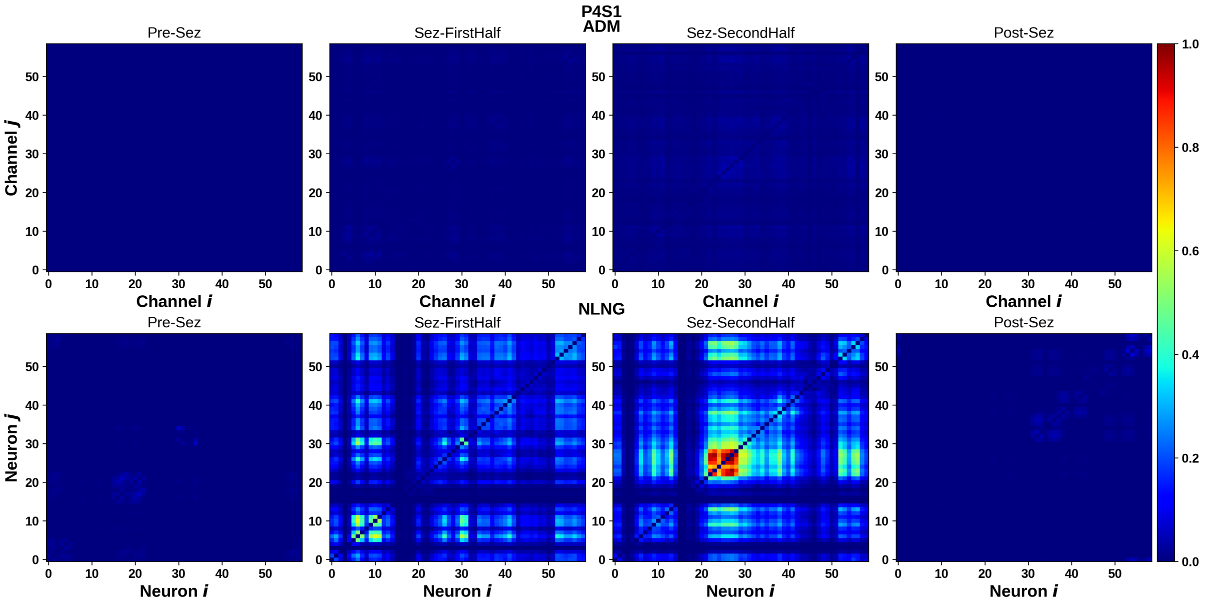


(a) Patient 4, Seizure 1

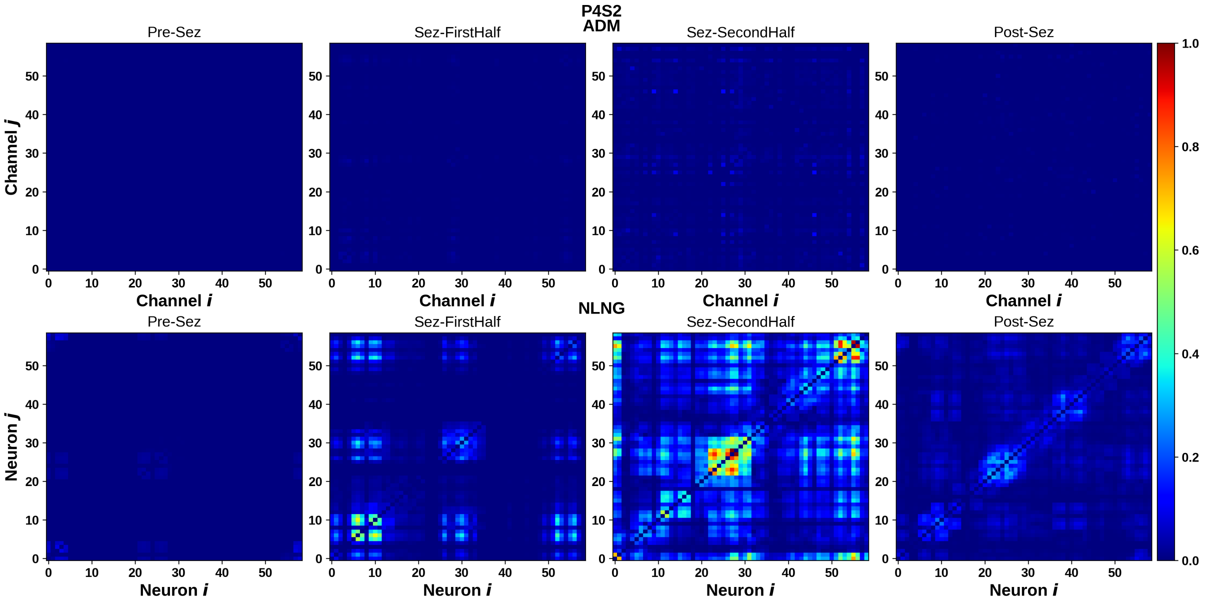


(b) Patient 4, Seizure 2

Supplementary Figure SF14: Synchronization matrices of Patient 4. The top row depicts the ADM/AFE channels and the bottom row depicts the NLNG neurons. The columns from left to right show synchronization during the pre-Seizure, first and second half of the Seizure, and post-Seizure periods respectively.

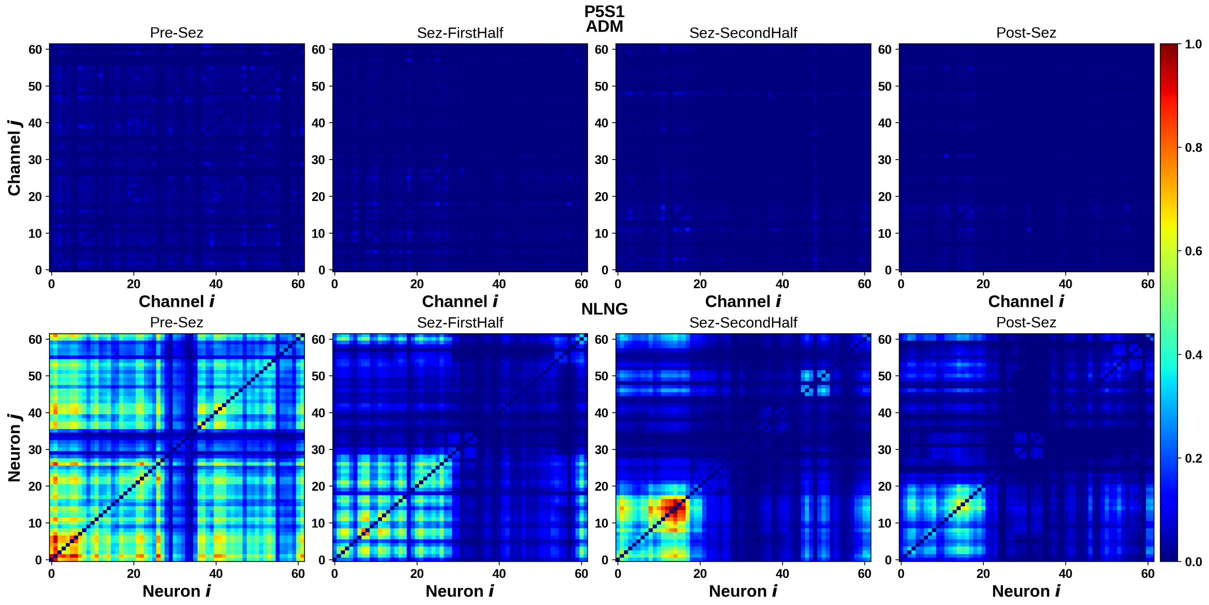


(a) Patient 5, Seizure 1

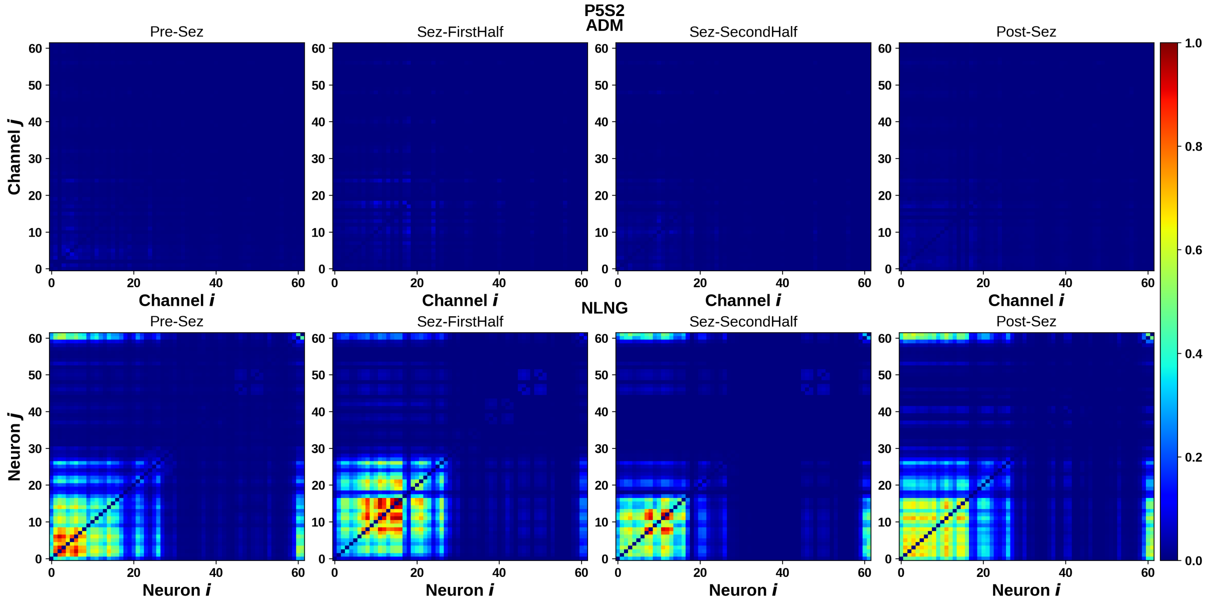


(b) Patient 5, Seizure 2

Supplementary Figure SF15: Synchronization matrices of Patient 5. The top row depicts the ADM/AFE channels and the bottom row depicts the NLNG neurons. The columns from left to right show synchronization during the pre-Seizure, first and second half of the Seizure, and post-Seizure periods respectively.

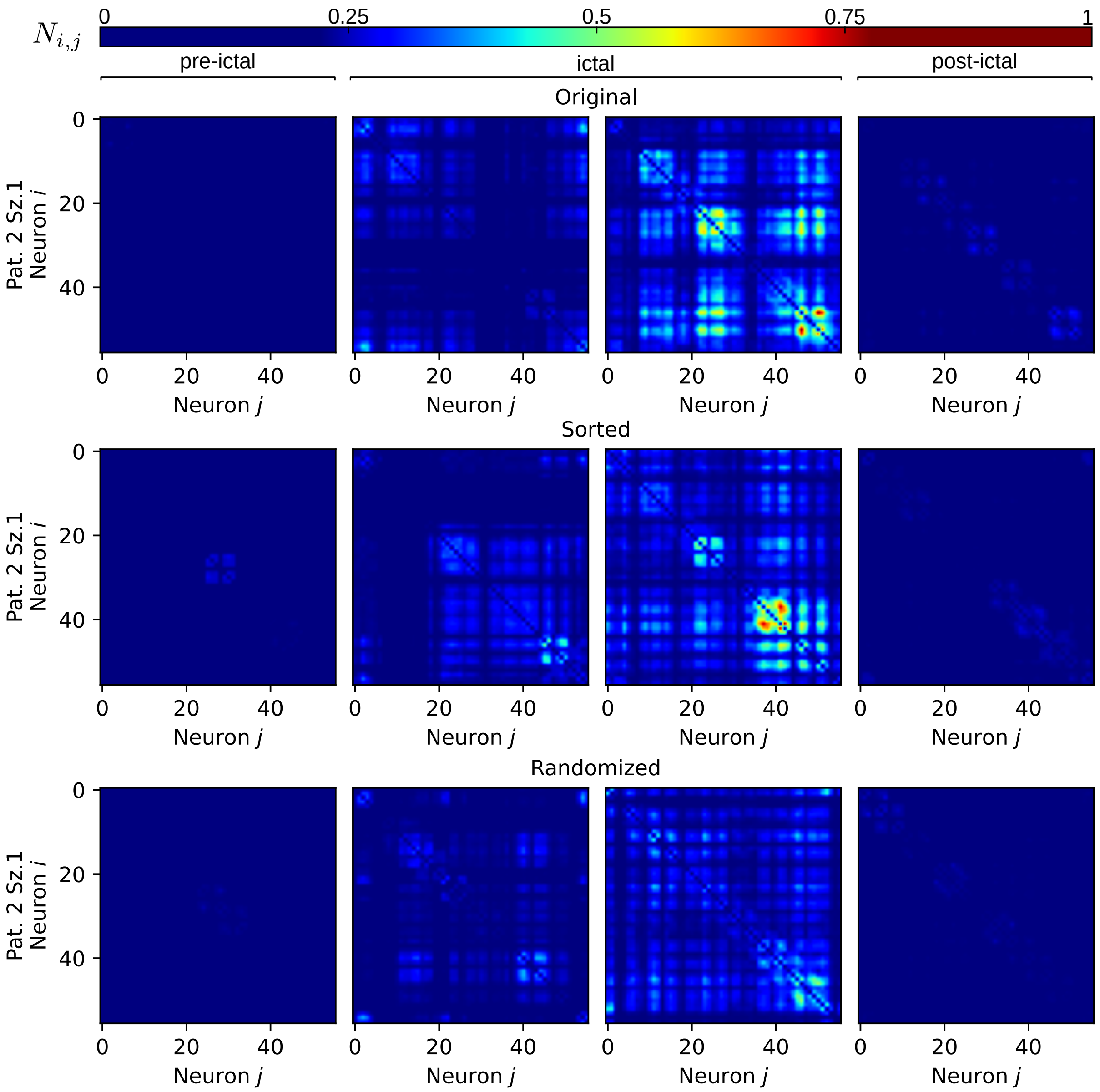


Supplementary Figure SF16: Synchronization matrices of NLNG neurons for two seizures of patient 2 seizure 1 based on three distinct input channel ordering methods. The first, second, and third rows represent synchronization based on input with the original dataset order, maximum correlation of the original data, and randomized order respectively.

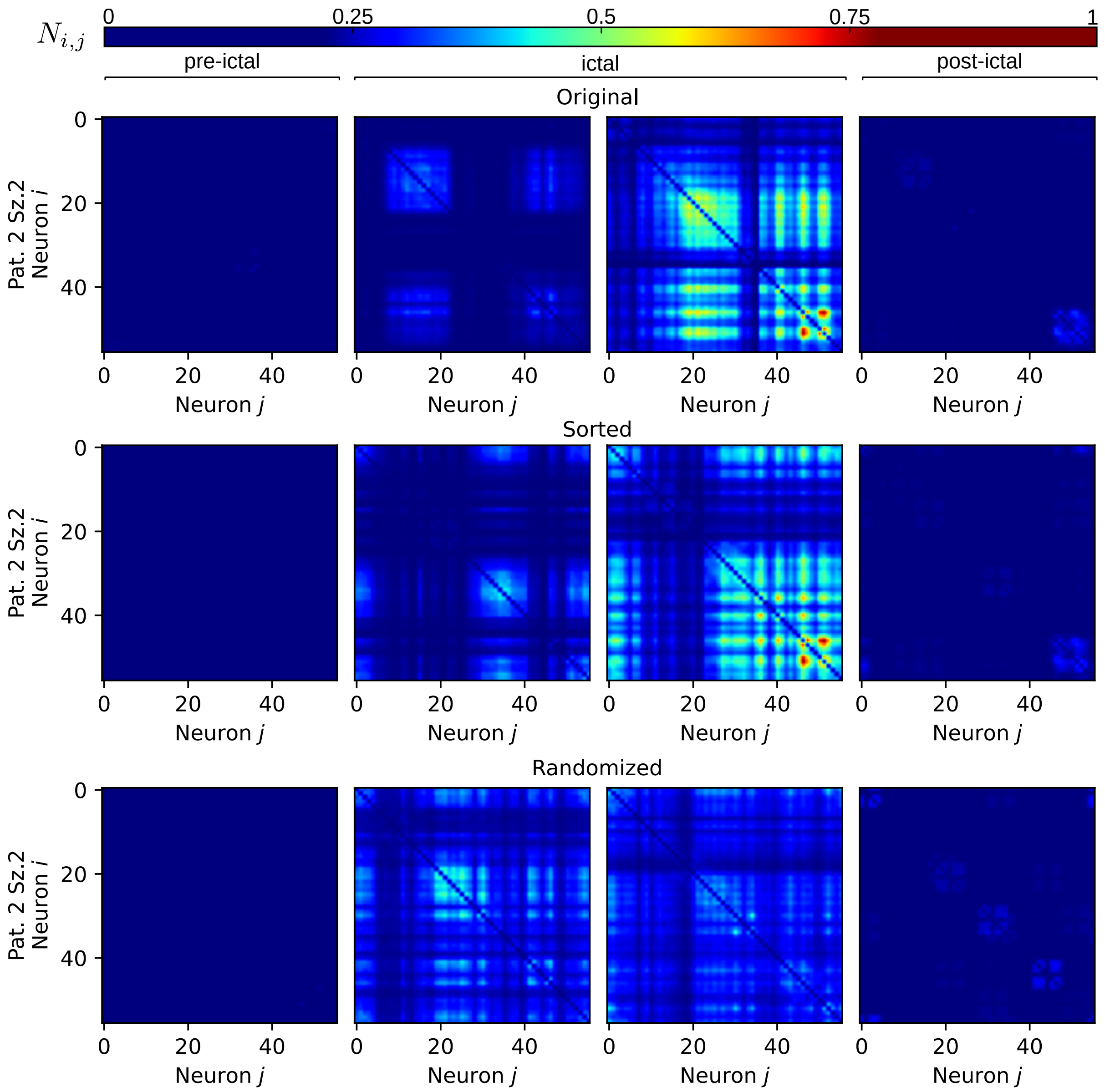


Supplementary Figure SF17: Synchronization matrices of NLNG neurons for two seizures of patient 2 seizure 2 based on three distinct input channel ordering methods. The first, second, and third rows represent synchronization based on input with the original dataset order, maximum correlation of the original data and randomized order respectively.

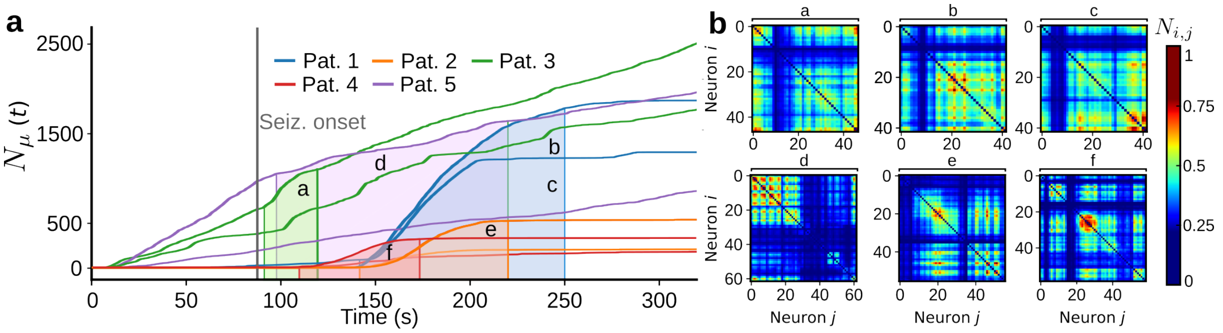


Supplementary Figure SF18: **a** The mean synchronization over all pairs of the NLNG layer cumulatively summed over time, colored areas under the curve indicate selected periods of time used to construct synchronization matrices depicted in **b**; Note two seizures are analysed for each patient and they correspond to two curves of identical color depicting the same. **b** Synchronization matrices taken during periods of maximal synchronization.

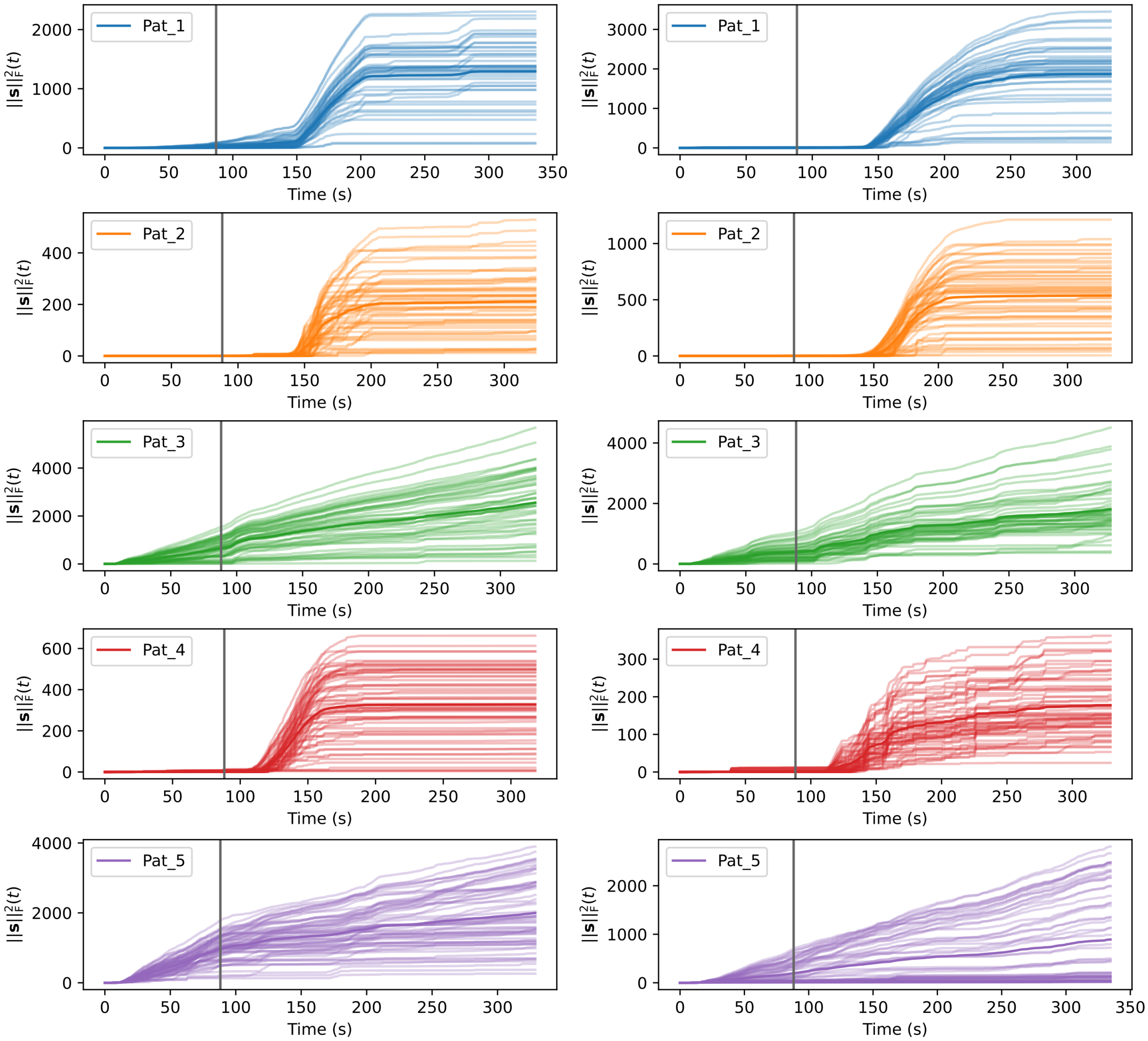


Supplementary Figure SF19: The synchronization across all non-local non-global layer neuron pairs over time for all patients and seizures. The grey line indicates the seizure start.

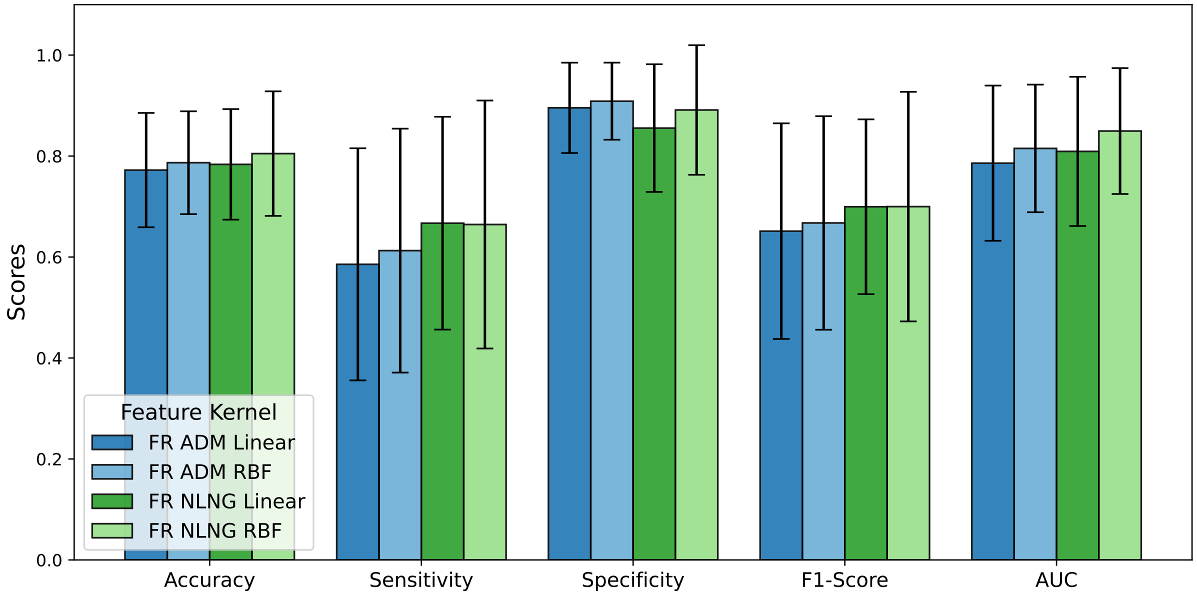


Supplementary Figure SF20: Averaged performance metrics across patients using SVM with Linear and RBF kernels based on the firing rates for ADM/AFE input and NLNG output.

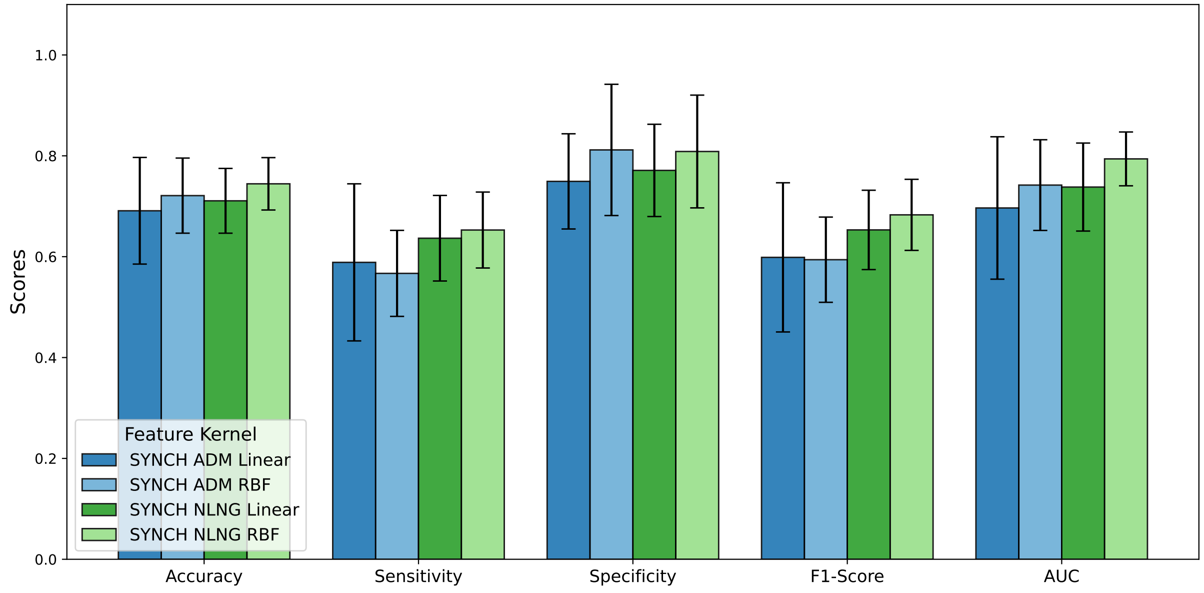


Supplementary Figure SF21: Averaged performance metrics across patients using SVM with Linear and RBF kernels based on synchronization measure for ADM/AFE input and NLNG output.

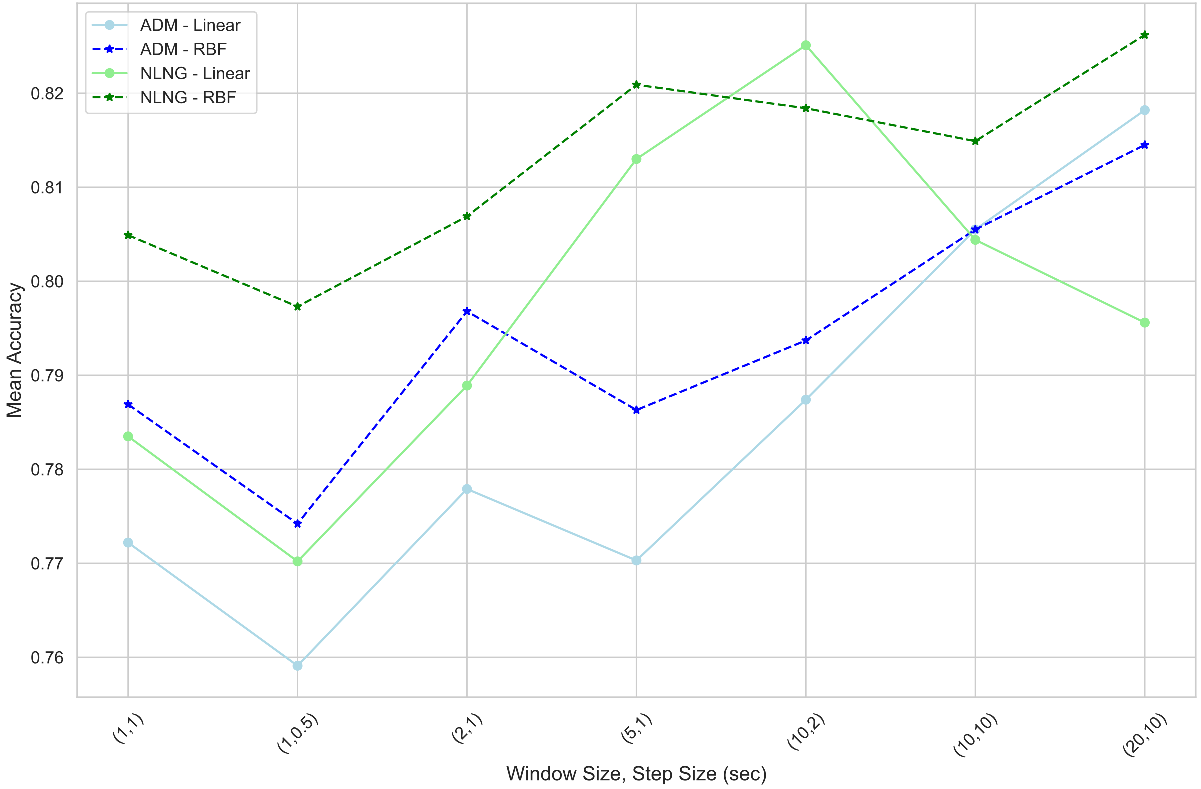


Supplementary Figure SF22: Accuracy across step and window sizes of the firing rates using SVM with Linear and RBF kernels for ADM/AFE input and NLNG output
